# Contribution of the lateral extension of the hyporheic zone in gravel bars in shaping river invertebrate diversity

**DOI:** 10.1101/2025.08.02.668304

**Authors:** Mc Jervis S. Villaruel, Junjiro N. Negishi, Junyi Wu, Shuyu Yao, Janine T. Mihara

## Abstract

Gravel bars (GBs) and hyporheic zone (HZ) are two major natural geomorphic features in gravel-bed rivers, having multiple functions ranging from habitat provisioning to nutrient cycling. This study examines whether the lateral extension of HZ in GBs contributes to the total diversity of river aquatic invertebrates. The hyporheic invertebrate community composition was entirely dissimilar from the benthic habitat, with stable GB hyporheic location exhibiting a more distinct community composition compared to wetted channel and active GB across all seasons. The local contribution to beta diversity (LCBD) also showed that stable GB has significantly higher uniqueness across all seasons, indicating a more unique invertebrate community. Species contribution to beta diversity (SCBD) indicates that this uniqueness was mainly driven by hyporheic dwelling species amphipod, *Pseudocrangonyx yezonis*, and Isopod species. Distance from wetted channel and dissolved oxygen were the factors influencing hyporheic invertebrates distribution. These findings show that the maintenance and restoration of gravel bars are crucial to enhancing and maintaining the overall invertebrate diversity in the river ecosystem.

## Introduction

Surface and subsurface features (i.e. landforms and alluvial sedimentation depth) of natural rivers are formed by geomorphic processes including constant erosion, transport, and sediment deposition, providing a critical foundation for various ecosystem structures and functions. Gravel bars (GBs) and hyporheic zone (HZ) are two well-known geomorphic features with high ecological importance in gravel-bed rivers. GBs are naturally occurring sedimentary formations formed by the deposition of gravel-sized particles through fluvial processes (Ock et al. 2015; Boodoo et al. 2017). GBs directly provide habitats for a wide variety of terrestrial flora and fauna (Kollmann et al. 1999; Zeng et al. 2015; Harvdova et al. 2023) and indirectly form optimal spawning grounds for fish (Geist and Dauble 1998; Beechie et al. 2010). The hyporheic zone (HZ), an active ecotone between surface water and groundwater in saturated sediment (Boulton et al. 2010; Mugnai et al. 2015), extends through the subsurface domain of wetted channels and gravel bars (Hauer et al. 2016). HZ is vital for nutrient cycling, organic matter processing, and water quality regulation (Boulton et al. 2010; Peyrard et al. 2011). In addition, the HZ provides a habitat for diverse biological assemblages (Lawrence et al. 2013; Stubbington et al. 2016; Datry et al. 2017). However, despite these conceptually recognized functions, knowledge of the quantitative importance of HZ in shaping invertebrate diversity in gravel-bed rivers remains limited.

Hyporheic invertebrates constitute a diverse assemblage of fauna that includes insects, crustaceans, and annelids, and perform critical ecological functions such as facilitation of nutrient cycling through the decomposition of organic matter, sediment processing through bioturbation, and energy transfer by serving as food resource for predators (Boulton et al. 2010; Lawrence et al. 2013; Stubbington et al. 2016; Datry et al. 2017). Diversity in hyporheic communities is determined by environmental filtering of species in relation to spatial gradients in water quality and sediment features along vertical and lateral dimensions of HZs (Peralta-Maraver 2018) and variable use modes of HZs. For example, some hyporheic taxa are adapted to low hyporheic DO (Malison et al. 2020) and limited spaces (Marmonier et al. 1993; Descloux et al. 2014). Furthermore, species with amphibiotic life cycles, such as insects with terrestrial adult stages, utilize the HZ only for larval development (Dole-Olivier et al. 2022). The HZ also provides a stable refuge, shielding organisms from natural disturbances, including flooding, predation, and extreme temperature fluctuations (Boulton et al. 2004; Davy-Bowker et al. 2006; McGrath et al. 2007; Robson et al. 2011). These temporal uses may occur largely in HZs with proximity to wetted channels. In contrast, certain amphipods and isopods species, spend their entire life cycle within the HZ as permanent inhabitants (stygobionts or permanent hyporheos) (Marmonier et al. 1993; Williams et al. 2010; Hutchins et al. 2020; Dole-Olivier et al. 2022). Thus, HZs in GBs may play significant roles in shaping invertebrate diversity by providing a wide range of the above-mentioned environmental gradients in the lateral dimensions, however its quantitative importance is unknown.

Beta diversity, the variation in species composition between different habitats, locations, or time points, quantifies how distinct or similar communities are across spatial or temporal scales (Whittaker 1972; Tuomisto, 2003). Beta diversity metrics provide valuable insights into the changes in community composition and structural patterns along environmental, spatial, and temporal gradients, and it is essential for understanding the mechanisms that influence species distribution (Jankowski et al. 2005; Anderson et al. 2011; Coelho et al. 2018). Legendre and De Cáceres (2013) introduced a method for partitioning β-diversity to assess the contributions of individual sites and species to overall community variation. This approach is represented by two metrics, the local contribution to β-diversity (LCBD), which quantifies the ecological uniqueness of a site based on species composition, and the species contribution to β-diversity (SCBD), which measures the relative importance of each species in shaping β-diversity patterns (Heino and Grönroos 2017; Siegloch et al. 2018). LCBD and SCBD have been widely applied in ecological studies, including assessments of species distribution shifts in diatom communities (Jyrkänkallio-Mikkola et al. 2018), fish assemblages (Gavioli et al. 2019), and macroinvertebrate communities (Heino and Grönroos 2017). These metrics can also provide direct and useful applications for river biomonitoring and management at a regional scale as well as informative indicators to identify sites to be protected to preserve riverine biodiversity (Ruhi et al. 2017; Tolonen et al. 2018; Li et al. 2020). Although beta diversity contribution assessments have been applied to hyporheic communities in wetted channels (Stubbington et al. 2015; Stubbington et al. 2019), no previous studies have applied this framework to the lateral extension of HZs beneath GBs.

This study examines whether the lateral extension of HZ in gravel bars (GBs) contributes to the total diversity of river aquatic invertebrates across different seasons. Specifically, we aim to (1) compare the invertebrate community compositional differences between benthic and various hyporheic locations (wetted channel, active GBs, and stable GBs), (2) evaluate the uniqueness of invertebrate communities within these habitats and identify the taxa driving this uniqueness, and (3) determine the environmental factors that best explain variations of invertebrate community across hyporheic locations. We hypothesize that the lateral extension of the HZ within GBs will consistently contribute to total invertebrate diversity across all seasons by harboring taxa unique to the hyporheic environment. We predict that permanently dwelling HZ taxa, such as amphipods and isopods species, occur disproportionately more in HZs beneath GBs and that their occurrence is associated with environmental gradients in DO and distance from the wetted channels. Previous studies demonstrated that unique species occur in GBs relative to wetted channel HZs (Mori and Brancelj 2011; Dole-Olivier et al. 2022). By quantifying the ecological role of GBs in enhancing invertebrate diversity, our study could provide a foundation for restoring and maintaining GBs as well as the geomorphic processes forming them especially in the face of escalating threats posed by river regulation activities.

## Methods

### Study site

The study was conducted between June 2022 to June 2024 in the Satsunai River, which spans a catchment area of 725 km^2^ with a channel length of 82 km. This river originates from Mt. Satsunai (42°41 N, 142°47 E; 1,895 m above sea level) in the Hidaka Mountain Range and flows to the Tokachi River in Hokkaido Prefecture, Japan. The mean daily discharge of the Satsunai River, measured at the Kami-Satsunai observation station (MLIT) located 12 km upstream of the study reach, was 12.9 m³/s over the period from 2010 to 2020.

The socio-economic need to deepen GBs ecological function is high in the Satsunai River. The size and formation of gravel bars have significantly diminished due to the accelerated forestation within the river channel. This transformation is primarily attributed to a substantial reduction in annual maximum river discharge, driven by the construction of the Satsunai Dam and other contributing factors (Takahashi and Nakamura 2011). Before the dam’s construction in 1998, the Satsunai River was a bar-braided, gravel-bed system characterized by extensive gravel bar formations (Sumitomo et al. 2018; Nakamura et al. 2020). To address this ecological issue, the Ministry of Land, Infrastructure, Transport, and Tourism (MLIT) initiated a restoration project in 2012 aimed at revitalizing gravel bars and riparian habitats. As part of this effort, an artificial flood was implemented by releasing a peak water discharge of 120 m³/s at the end of June, a magnitude equivalent to the two-year return period flood before the dam’s construction (Nakamura and Shin 2001).

Two sites were selected, one site in the upstream (L7) and downstream area (L2) (Fig. 1). The hyporheic pipe and installation design was adapted from the study of Negishi (2023). In total, 15 hyporheic pipes were installed per site on June 9, 2022 (total of 30), in a gravel bar to collect the hyporheic invertebrate communities and measure environmental conditions. The pipes were made of PVC pipes (10–15 cm in diameter) with lower-end openings covered by either polyethylene or stainless-steel mesh screens (mesh size of 4 mm × 3 mm) (Supplementary file 1 Fig. S1) over a vertical distance of 20 cm at the lowest end. Three hyporheic locations with varying horizontal distances from the wetted channel were considered during pipe installation to capture the lateral extension of HZ in the GBs. These locations include the wetted channel (T1: situated within the wetted channel), active gravel bar (T2: horizontal distance from wetted channel 10.8-38.9 m), and stable gravel bar (T3: horizontal distance wetted channel 40.5–83.1 m) (Fig. 1; Supplementary file 1 Fig. S2). During periods of flow recession after the installations, drying of the T1 location exposed some pipes on the dry surface, resulting in a maximum distance of up to 1.9 m from the base flow. The distances of each pipe location varied temporarily because the pipe locations changed over time as floods caused damages and changes in landforms. A new set of T1 and T2 pipes was installed in a point bar in L2 in November 2022 (Supplementary file 1 Fig. S2 C). These newly installed pipes, along with the remaining T3 pipes, were utilized for the subsequent field collections and observation. A new set of pipes was installed in L2 in May 2024, in addition to the existing old T2 pipes. At the same time, an additional set of hyporheic pipes was also installed on an island bar (IB) near the L2 site (Supplementary file 1 Fig. S2 D). The elevation of ground surfaces relative to the river surface during base-flow conditions varied among the three hyporheic locations with <19 cm elevation for T1, 25–98 cm for T2, and 105–152 cm for T3. Despite these differences, it was ensured that at least 50 to 60 cm of the lower end of the pipe remained submerged within the groundwater zone. This is to ensure that the groundwater levels across all hyporheic pipes were comparable and closely matched the elevation of surface flow within the river channel.

**Fig. 1.**
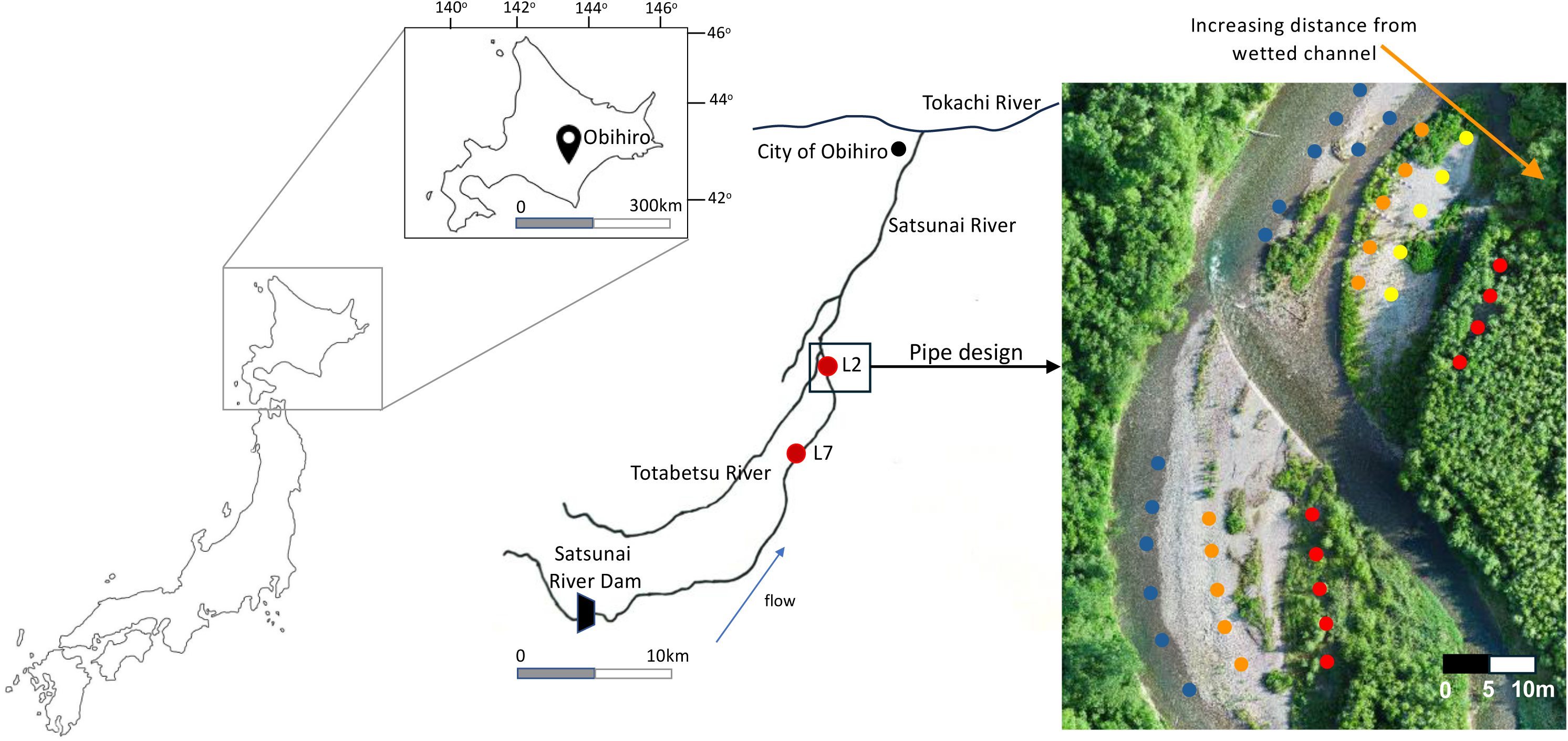
Location of the study site. The red dot in the map shows the sampling points: L7 - upstream; and L2 - downstream. The L2 site on June 17, 2024, is displayed as an example in the rightmost part, highlighting the pipe design with increasing distance from the wetted channel (blue = wetted channel; orange and yellow = active gravel bar; red = stable gravel bars).

### Invertebrate sample collection

Benthic and hyporheic invertebrates were collected on five occasions covering various seasons. Collections were done on June 19-20, 2022 (early summer), December 6, 2022 (winter), March 29, 2023 (spring), May 25, 2023 (late spring), and June 24, 2023 (early summer). Preceding the collection of hyporheic colonization traps, benthic invertebrates were collected using a Surber sampler (0.0625 m^2^, 324-µm mesh) (total of 35 benthic samples; Supplementary file 2 Table S1). To ensure concordance with the hyporheic community, benthic samples were specifically collected from the closest wetted channel associated with each T1 pipe. The collection of benthic samples was done starting from the downstream-most area of each site, totaling 5 samples per site.

Hyporheic invertebrates were collected using hyporheic colonization traps (total of 24 hyporheic samples from T1, 24 from T2, and 17 from T3; Supplementary file 2 Table S1). Colonization traps were made following the procedure of Negishi et al. (2019). Hyporheic colonization traps were made using the available riverbed sediments in the study site. The sediments collected in situ were washed thoroughly to remove detritus and invertebrates. For 10-cm pipes, 707 cm^3^ sediments (D_50_ = 8–16 mm) were tightly packed in a polyethylene mesh (4 × 5 mm mesh size). Dried Japanese Alder leaves (*Alnus japonica*) weighing 3 (±0.05) grams were also placed along with the sediment to mimic organic matter contents in the sediment. The trap was further encased in a durable mesh casing (mesh size = 3 mm × 3 mm) to ensure that the sediments were secure (Supplementary file 1 Fig. S3). For 15-cm diameter pipes, 3004 cm^3^ sediments (D_50_ = 8–6 mm) were tightly enclosed in a 3 × 4 mm mesh bag. A rope was attached to each trap and installed inside each hyporheic pipe. Moreover, temperature loggers (HOBO 8K Pendant; Onset Co., Bourne Massachusetts) and plastic tubes (an internal diameter of 6 mm) were installed inside the pipe alongside each trap, however, loggers were not installed to all pipes due to limited availability. The plastic tubes served as conduits for the collection of water samples used in nutrient analysis and water quality assessments. All pipes were covered with a PVC lid cover after trap installation. Hyporheic traps were collected after an incubation period of 13 to 60 days to ensure adequate time for invertebrate colonization.

Hyporheic invertebrates were collected by retrieving colonization traps installed within the hyporheic pipes. The traps were accessed by removing the PVC lid with a hammer and swiftly pulling the attached rope to retrieve the traps. The traps were immediately caught by a hand-held D-frame net (375 µm) to ensure that the invertebrates that may fall from the traps could be caught. Both benthic and hyporheic samples underwent rinsing to eliminate sediments and organic matter before sieving through a 500-µm stainless-steel mesh. The invertebrate samples were preserved using 75% ethanol and transferred to the laboratory for further analysis. Invertebrates were then sorted and counted under a stereomicroscope and were identified to the lowest possible taxonomic level using the taxonomic keys of Kawai and Tanida (2005) and Merritt et al. (2019).

### Measurement of environmental parameters

Measurements of water quality parameters were done for both hyporheic and benthic sites. Hyporheic water samples were obtained by manually pumping water from plastic tubes installed within each hyporheic pipe using a manually operated bilge pump. Subsequently, the withdrawn hyporheic water was utilized for the point measurement of dissolved oxygen (DO) and temperature using a portable oxygen meter (HQ40d portable DO meter, Hach Ultra Co., Tokyo, Japan). Electrical conductivity (EC) and pH were also measured using a hand-held probe (WM 32EP, DKK-TOA Co., Tokyo, Japan). Water quality measurements for the benthic habitat were taken directly from the surface water in the locations where benthic invertebrate samples were collected. In addition to these parameters, water depth and 60%-depth velocity were measured three times for each benthic sample. Additionally, hyporheic and benthic water samples (50 mL) were also collected using an acid-washed falcon tube for the analysis of total nitrogen (TN), total phosphorus (TP), and concentrations of specific ions (Cl^-^, PO_4_^2-^, F, NO^2-^, Br, NO_3_^2-^, and SO_4_^2-^). Following collection, the samples were stored in a cooler box, and transported to the laboratory for subsequent analysis. TN and TP concentrations were determined by digesting unfiltered water samples using the colorimetric method outlined by Rahman et al. (2021). The absorbance of the samples was subsequently measured using a spectrophotometer (UV-1280, Shimadzu Co., Japan). For the ion analysis, water samples were first filtered using a glass fiber filter with a pore size of 0.45 µm (GC-50, Advantec Co., Tokyo, Japan) before running in an ion analyzer-chromatograph (IA-300, TOA-DKK Co., Tokyo, Japan).

The exact distance of each pipe from the wetted channel was determined using QGIS (version 3.40; QGIS Development Team 2024). GPS coordinates for each pipe installed in L2 and L7 during the 2022 and 2023 field collections were recorded and plotted in Google Earth Pro to ascertain their geographical positions. This georeferenced map was imported to QGIS. For the June 2024 collection, high-resolution aerial imagery of the pipe locations was captured using an unmanned aerial vehicle (UAV) (DJI Mavic 3M and DJI Matrice 300 RTK with DJI Zenmuse L1 at altitudes of 150 m and 70 m, respectively). The UAV imagery was imported to QGIS to accurately map the pipe locations in the L2 and the IB sites. The geographic positions of individual pipes were determined based on visible markings. The “measure line” tool in QGIS was subsequently used to determine the shortest linear distance from each pipe to the nearest point on the wetted channel.

### Statistical analyses

The differences in the invertebrate community composition among locations over different seasons were examined visually via non-metric multidimensional scaling (NMDS) using the Bray-Curtis similarity matrix. The statistical differences associated with the seasons, locations (benthic and various hyporheic habitats), and their interaction were assessed using a 2-way permutational multivariate analysis of variance (PERMANOVA) with 999 permutations. Where PERMANOVA showed significant differences, pairwise PERMANOVA was employed to scrutinize the differences between groups (Martinez-Arbizu 2019).

Taxonomic richness (α diversity) across different seasons was evaluated and compared between benthic and multiple hyporheic habitats. A generalized Linear Mixed Model (GLMM) was employed to investigate whether taxonomic richness exhibited variation among various locations across different seasons. In this model, taxonomic richness was designated as the response variable, with season, location, and their interactions as the main factors, while the site was included as a random factor (Poisson error distribution). Similarly, abundance was compared between benthic and various hyporheic habitats. GLMM was utilized to examine whether abundance fluctuated across locations and seasons. Abundance was set as the response variable, with season, location, and their interactions as the main factors, while the site was incorporated as a random factor (negative binomial error distribution). The relative abundance of each invertebrate taxa in benthic and hyporheic locations over various seasons was also calculated to identify the most abundant taxa in each location. The relative abundance (%) of species “a” was calculated separately for each location (benthic, T1, T2, T3) during each seasonal collection by dividing the abundance of species “a” by the total abundance of all species within that specific location.

The uniqueness of the invertebrate community across different locations and seasons was assessed using local contribution to β-diversity (LCBD). The LCBD was calculated for each sample collected in different locations in each sites using the equation: LCBDi = SS_i_/SS_total_, where SS_i_ is the squared distance of the sampling site i to the average site (centroid) in the multivariate ordination distance space, and SS_total_ is the sum of all SS_i_ values across sites (Legendre and De Cáceres, 2013). To dampen the influence of highly abundant species, community abundance data were subjected to Hellinger transformation prior to analysis. This transformation increases the sensitivity of LCBD to the presence and relative proportions of taxa, rather than relying solely on raw abundance counts (Legendre and Gallagher 2001; Legendre and Legendre 2012; Legendre and De Cáceres 2013). LCBD values were then compared across habitat locations and seasons using GLMM. In this model, LCBD values were treated as the response variable, with seasons, sampling locations, and their interactions as the main predictors. A Gaussian error distribution was applied, with the site included as a random effect. Additionally, species contribution to β-diversity (SCBD) was calculated for each species using the Hellinger-transformed abundance data to identify which taxa drive the observed uniqueness among the different locations. SCBD was calculated using the equation: SCBD_j_ = SS_j_/SS_total_ where SS_j_ is the contribution of species j to the overall β-diversity, calculated as the sum of the central and square values of species j, and SS_total_ is the sum of all SS_j_ values across all species (Legendre and Cáceres 2013).

Redundancy Analysis (RDA) was conducted on the Hellinger-transformed data to identify the environmental parameters that influenced the hyporheic invertebrate community (Legendre and Legendre 2012). First, the degree of multicollinearity among the 10 environmental parameters was examined using variance inflation factors (VIF). If any VIF exceeded 4, forward selection was used to iteratively remove redundant variables until all remaining variables had VIFs below 4 (Hu et al. 2022). The significance of the RDA model was also assessed with 999 permutations. In addition, PERMANOVA was employed to test whether there is a difference in centroids of groups (hyporheic locations) in the RDA multivariate space based on RDA scores, utilizing the Bray-Curtis similarity index, with significance determined via 999 permutations. Subsequently, pairwise PERMANOVA was performed to compare different locations. This analysis allowed us to identify which groups exhibited differences in their community composition in response to the environmental parameters.

The relationship between thermal stability and dissolved oxygen (DO) in the hyporheic habitat was investigated using water temperature data collected by the temperature loggers. This separate analysis was done because logger data was limited, making it impossible to include it in the RDA analysis. Thermal stability at each site was quantified by calculating the relative coefficient of variation (CV) of hourly water temperature measurements over a four-day period prior to sampling, serving as an indicator of temperature fluctuations and stability. Non-linear logarithmic regression analysis was conducted to test their relationship and to provide insights into the interplay between temperature dynamics and oxygen availability in the different hyporheic locations.

All statistical analyses (significance threshold = 0.05) were performed using R (version 4.2.0; R Core Team 2023). The package “vegan” was used for NMDS and “adonis2” for PERMANOVA with pairwise.adonis2 function to check the pairwise comparison. For the LCBD and SCBD, the “adespatial” package was used (Legendre and De Caceres 2013). The RDA was conducted using the “vegan” package with the RDA model constructed using the rda function and anova.cca function to test whether the relationship between the environmental variables and community composition was statistically significant. The package “glmmTMB” was used for GLMM and the package “multcomp” for the multiple comparison.

## Results

The environmental parameters measured during the study is summarized in Table S2 of Supplementary file 2. The lowest electrical conductivity (EC) was recorded in the early summer of 2022 (3.60 mS/m) while the highest EC was noted in spring (6.72 mS/m). The coldest temperatures were observed in winter (2.44 °C), while the warmest occurred in late spring (17.58 °C) and early summer (17.51 °C). pH values remained near neutral across sites throughout the seasons, varying between 6.12 and 6.97. The highest dissolved oxygen (DO) concentration was noted in winter (11.74 mg/L; 87.80% saturation) whereas the lowest DO was recorded in summer (7.22 mg/L; 73.1% saturation). Total nitrogen (TN) reached its highest levels in spring (2.66 mg/L) while lowest in early summer of June 2022 (0.15 mg/L). Total phosphorus (TP) was consistently low across all locations throughout the study period (0.01 mg/L to 0.21 mg/L). Nitrate (NOLJ^2^LJ) exhibited its lowest concentrations in early summer (1.43 mg/L) and highest in spring (3.77 mg/L). The velocity of the surface water was 28.7 to 99.6 cm/sec while the shallowest depth was 25.8cm and the deepest depth was 37.7 cm.

A total of 14,241 individuals belonging to 71 invertebrate taxa were collected. This is distributed to 12,952 benthic individuals belonging to 66 taxa and 1,289 hyporheic individuals belonging 37 taxa (Table 1). Hyporheic community composition was entirely distinct from benthic habitat across all seasons based on the NMDS plot (stress=0.190; Fig. 2). The two-way PERMANOVA analysis indicated that community composition was significantly influenced by location (Table 2A). Subsequent pairwise comparisons revealed a marked differentiation between hyporheic and benthic communities. Within the hyporheic zones, sites T1 and T2 displayed relatively similar community compositions, whereas T3 exhibited a distinct and more differentiated assemblage compared to T1 and T2 (Table 2B). In addition, the benthic community demonstrated higher taxonomic richness and abundance than the hyporheic community (Supplementary file 1 Fig. S4 and S5; Supplementary file 2 Tables S3 and S4).

**Fig. 2.**
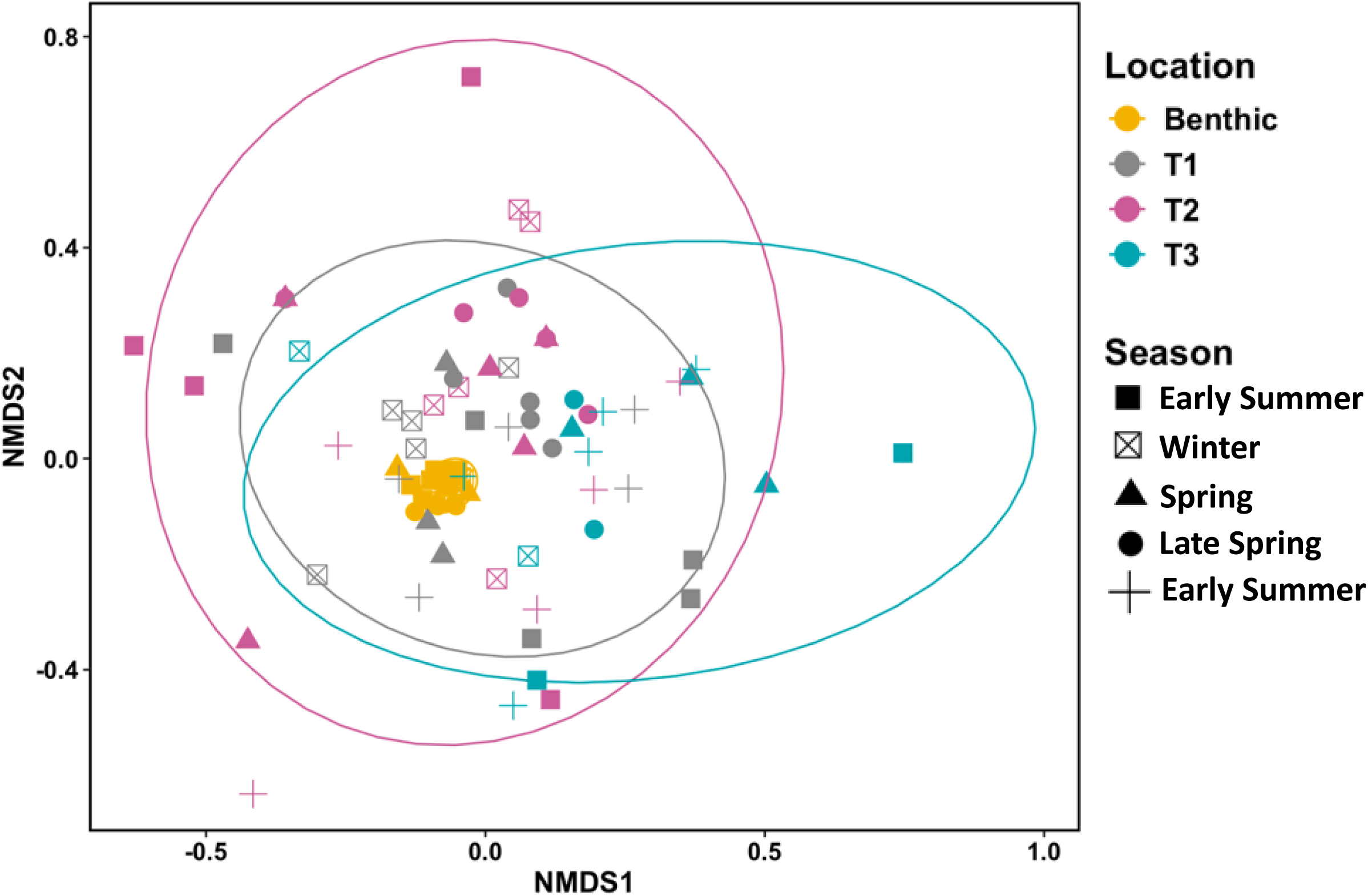
Results of nonmetric multidimensional scaling (NMDS), including 95% confidence interval ellipses for the invertebrate community composition dissimilarity between different locations over various seasons.

**Table 1.**
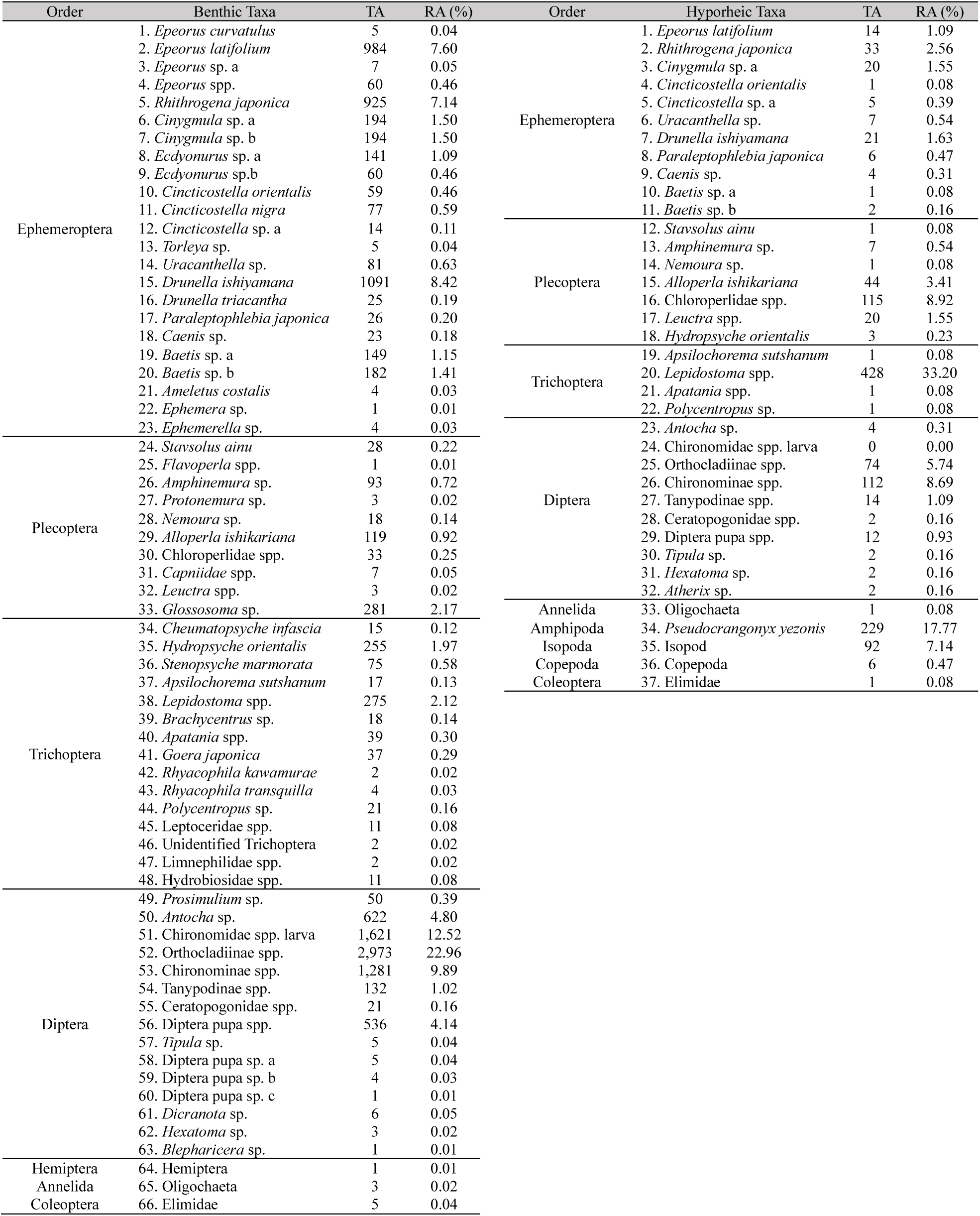
List of benthic and hyporheic invertebrate taxa, showing their total abundance (TA) and relative abundance (RA).

**Table 2.**
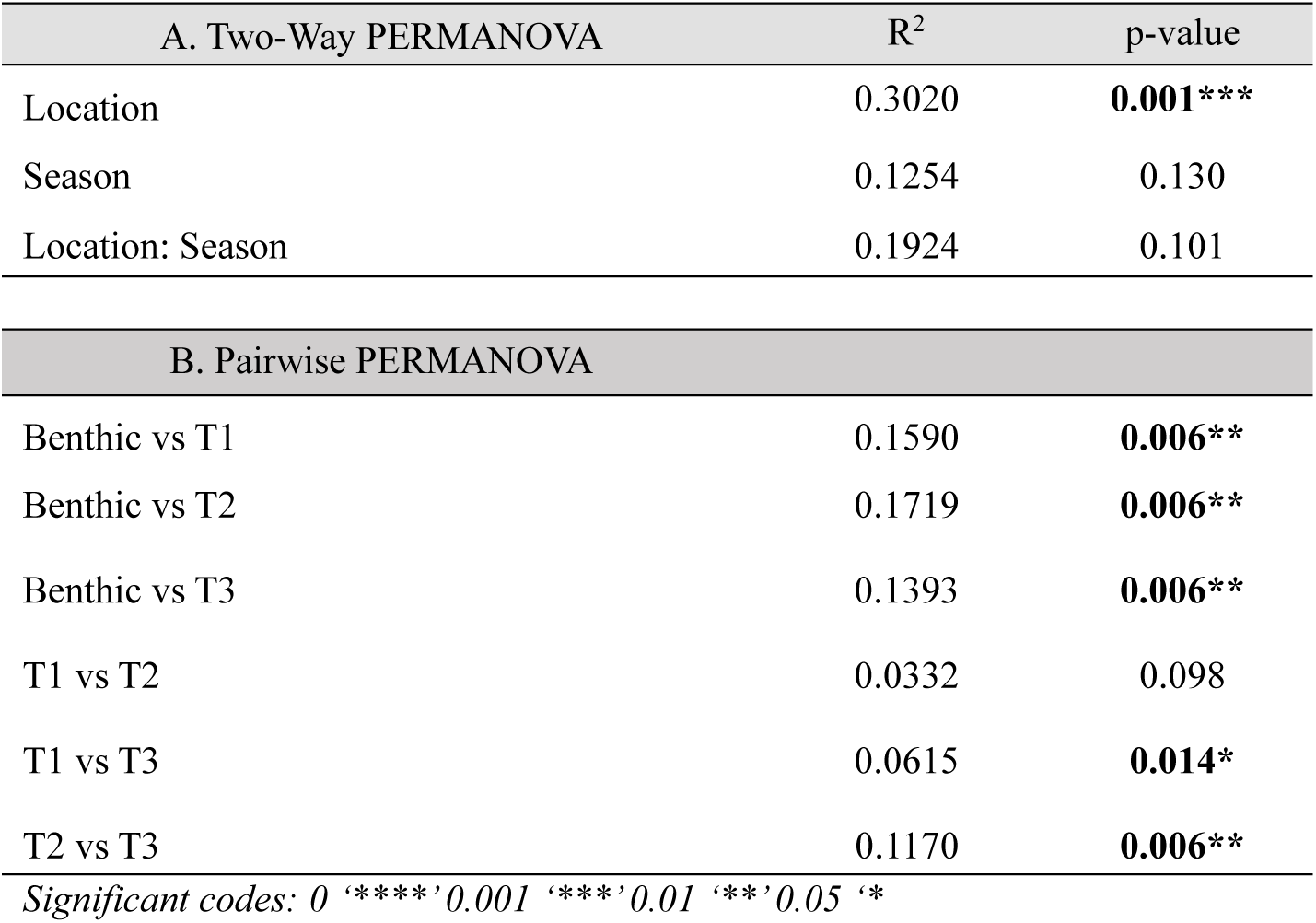
Results of the Two-Way Permutational Analysis of Variance (PERMANOVA) testing the difference of invertebrate community composition by location and season [A] and Pairwise PERMANOVA testing the difference of invertebrate community composition by location [B]. P-values in bold font indicate significance.

LCBD exhibited significant variation across locations (p<0.001). Multiple comparisons demonstrated that the T3 location consistently exhibited significantly higher LCBD values indicating a unique invertebrate community at T3 (Fig. 3; Supplementary file 2 Table S5). SCBD showed that various taxa contributed to the uniqueness of each location (Fig. 4). In the benthic location, *Rhithrogena japonica* consistently exhibited high SCBD values across four seasons at both L2 and L7 sites, indicating its strong contribution to the community uniqueness. Other Ephemeroptera taxa also showed elevated SCBD scores, although the specific contributing species varied among seasons and sites. Additionally, Diptera taxa, particularly members of the Orthocladiinae and Chironominae subfamily, consistently contributed to the uniqueness of benthic communities across different sites and seasons. T1 exhibited a distinct set of species influencing beta diversity, differing from those observed in the benthic habitat. Amphipoda species, *Pseudocrangonyx yezonis*, consistently contributed to the uniqueness of this location, as indicated by its high SCBD values across all sites and seasons. Similarly, the Isopoda species was also a key driver of the location’s uniqueness, exhibiting elevated SCBD values across three seasons at both the L2 and IB sites. Other taxa, including *Lepidostoma, Alloperla ishikariana, Leuctra*, Oligochaeta, and Copepoda, also consistently displayed higher SCBD values across sites, suggesting their significant role in shaping community uniqueness in T1. The SCBD values in the T2 locations similarly highlighted the same species as key contributors to beta diversity. *Pseudocrangonyx yezonis*, Isopoda, *Lepidostoma*, Oligochaeta, and members of the subfamily Chironominae consistently emerged as major contributors to the uniqueness of various sites across seasons. Additionally, Orthocladiinae species also demonstrated high SCBD values specifically in L2 and L7 sites in all seasons except in late spring. *Pseudocrangonyx yezonis* and Isopoda were noted to be key contributors to the uniqueness of T3 locations, having consistently high SCBD across all seasons and sites. Moreover, taxa such as *Lepidostoma, Alloperla ishikariana*, Oligochaeta, and Diptera larvae also displayed high SCBD values, particularly at the L2 site, underscoring their persistent role in structuring the uniqueness of T3 location. The mentioned taxa were also noted to be predominantly more abundant in the locations wherein they showed high SCBD values (Supplementary file 1 Fig. S6). For instance, *Pseudocrangonyx yezonis* and Isopoda not only exhibited consistently high SCBD values in T3 but were also predominantly abundant in T3 across all seasons.

**Fig. 3.**
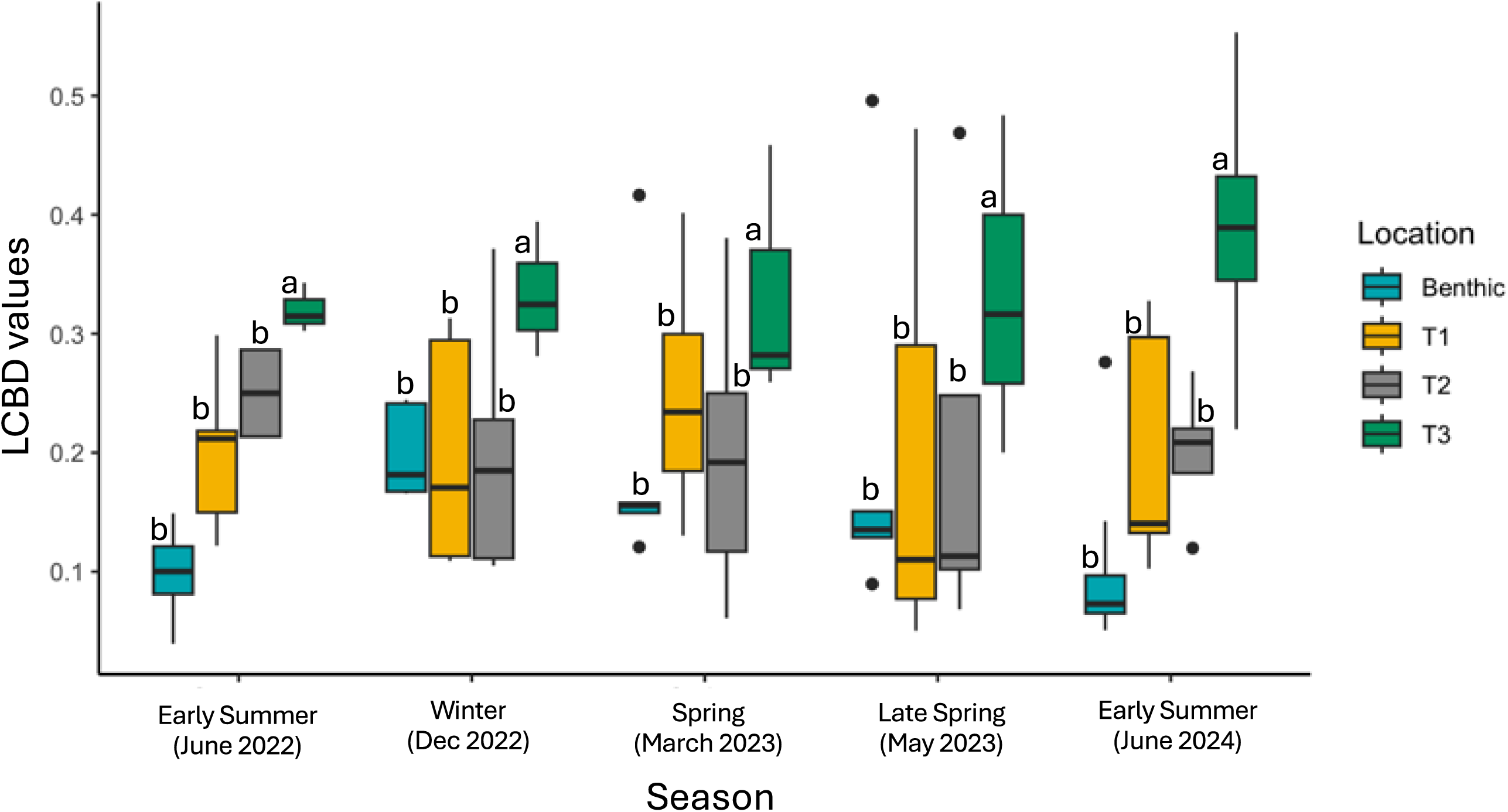
Comparison of Local Contribution to Beta Diversity (LCBD) values among different locations (benthic, T1, T2, T3) across seasons. Multiple comparison was conducted among locations only. Different letters indicate statistical differences based on multiple comparisons from the GLMM analysis; locations sharing the same letters are not significantly different.

**Fig. 4.**
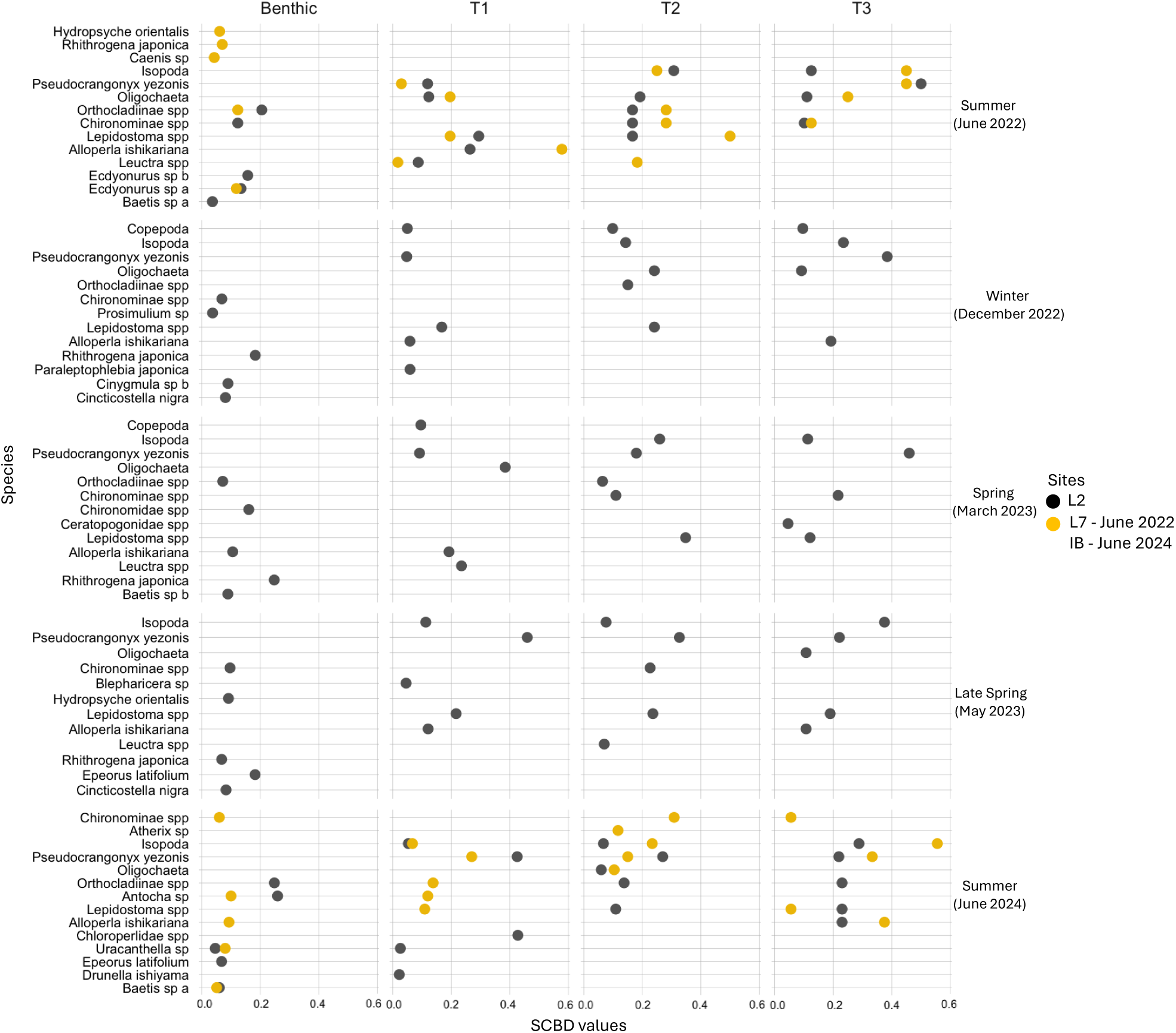
SCBD values of invertebrate taxa in benthic and various hyporheic locations across seasons and sites (L7, L2, and IB). SCBD scores are reported separately for each site, however, SCBD for winter, spring, and late spring seasons are presented only for the L2 site due to the loss of hyporheic pipes at L7 caused by flooding. Each circle represents the SCBD values of each taxa per site. The L2 site is represented in black, while L7 (early summer of June 2022) and IB (early summer of June 2024) are represented in orange. The top five taxa with the highest SCBD values are shown, except for T3 early summer (June 2022), where only four taxa were recorded.

The first two RDA axes explain a combined 14.27% of the total variance in community composition (axis 1 = 7.58%, axis 2 = 6.69%; F = 2.7339; p <0.001) (Fig. 5; Supplementary file 2 Table S6). Dissolved oxygen and distance from the wetted channel were the significant factors that influenced the separation of the hyporheic community in the RDA plot (Table 3). T1 was strongly associated with higher DO (p <0.01), with its points clustering along the DO vector in the upper right quadrant. In contrast, T2 and T3 hyporheic community composition was significantly associated with the distance from the wetted channel (p <0.01) showing a negative association. The PERMANOVA results confirmed a statistically significant difference among the groups in the RDA multivariate space (p <0.01) (Supplementary file 2 Table S7 A). Pairwise PERMANOVA analysis further indicated that the hyporheic invertebrate communities in gravel bars (T2 and T3) were significantly distinct from those in T1 (Supplementary file 2 Table S7 B), showing that environmental factors shaped hyporheic invertebrate community structure. Additionally, the temperature stability and dissolved oxygen (DO) showed a positive correlation (Fig. 6).

**Fig. 5.**
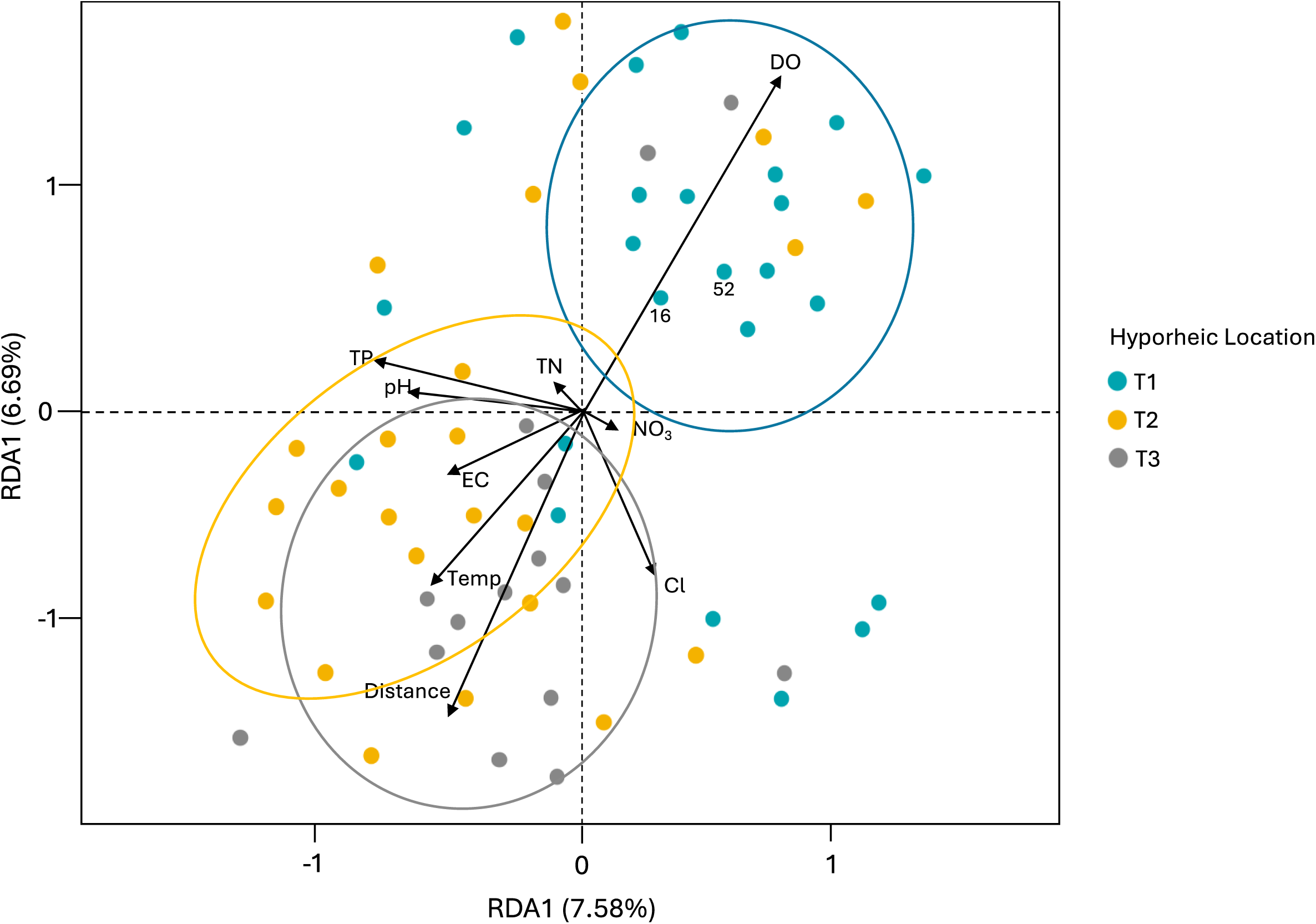
RDA ordination plots showing the association of the hyporheic community with the environmental parameters. Ellipses represent the 95% confidence intervals for each hyporheic location, showing the clustering of locations (denoted by colors).

**Fig. 6.**
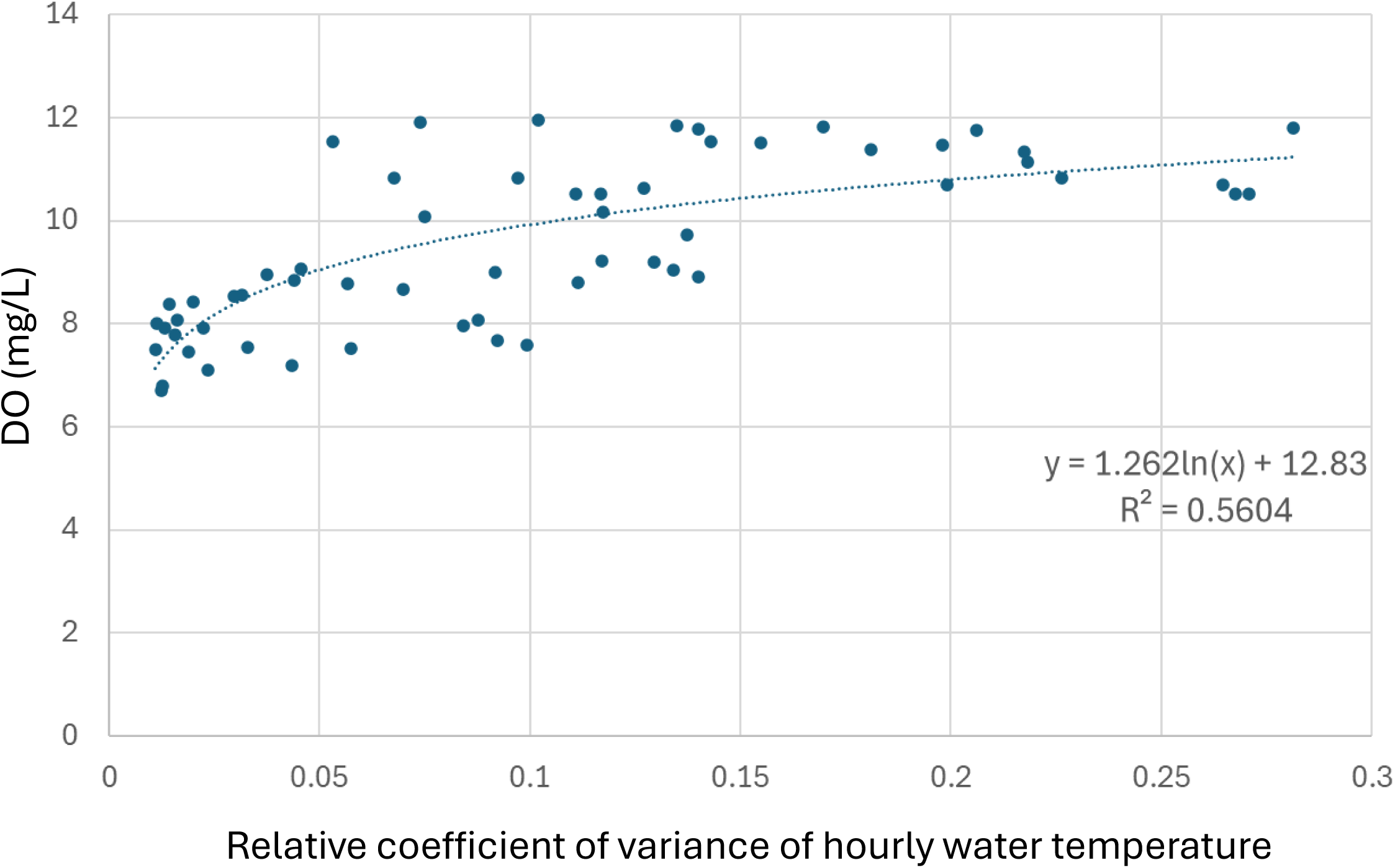
Non-linear logarithmic regression showing the relationship between the temperature locations and dissolved oxygen. The coefficient of variation (CV) of hourly water temperature measurements over a 4-day period prior to sampling was used as an indicator of thermal stability.

**Table 3.** Results of the Redundance Analysis (RDA). The bold lines are factors that showed significant correlation with hyporheic community. P-values in bold font signify significance.

|  | RDA1 | RDA2 | R<sup>2</sup> | Pr (> r) |
| --- | --- | --- | --- | --- |
| EC | -0.309 | -0.462 | 0.712 | 0.632 |
| temp | 0.121 | -0.353 | 0.625 | 0.132 |
| pH | -0.361 | -0.130 | 0.251 | 0.201 |
| DO | 0.028 | 0.646 | 0.181 | **0.002\*\*** |
| TN | -0.027 | 0.022 | 0.112 | 0.402 |
| TP | -0.547 | -0.147 | 0.009 | 0.550 |
| Cl | -0.088 | -0.326 | 0.005 | 0.080 |
| NO<sub>3</sub> | -0.020 | -0.013 | 0.003 | 0.402 |
| Distance | 0.355 | -0.707 | 0.019 | **0.006\*\*** |

Significant codes: 0 '\*\*\*\*' 0.001 '\*\*\*' 0.01 '\*\*' 0.05 '\*'

## Discussion

This study investigated whether and how the lateral extension of the hyporheic zone in gravel bars contributes to the total invertebrate diversity in rivers, and if the contribution varies seasonally. The study confirmed our hypothesis that the hyporheic habitats in gravel bars can contribute to the total river invertebrate diversity across various seasons. This contribution primarily stemmed from the high uniqueness of the stable gravel bar site (T3) which was mainly driven by hyporheic-dwelling taxa such as *Pseudocrangonyx yezoni*s and Isopod species. Although correlational approaches have limitations in elucidating causal relationships, environmental filtering by dissolved oxygen, thermal stability, and distance were found and suggested to be important. While previous studies have highlighted differences in invertebrate community composition and diversity between hyporheic and benthic zones, noting the presence of unique species in hyporheic habitat including those beneath gravel bars (De Walt and Stewart, 1995; Fowler and Death, 2001; Mori and Brancelj, 2011; Descloux et al. 2013; Marmonier et al. 2019; Negishi et al. 2019; Malison et al. 2020; Marmonier et al. 2020; Negishi et al. 2022), they did not elucidate the contribution of the lateral extension hyporheic habitat in the gravel bars to the overall invertebrate diversity in rivers and streams and thus lacked in temporal changes in contribution at seasonal time scales. Therefore, this study offers a novel perspective on the role and importance of hyporheic habitats in gravel bars in maintaining invertebrate diversity in rivers.

### Gravel-bar hyporheic zone as a driver of invertebrate diversity and their functions

Community composition in hyporheic zones was consistently distinct from benthic habitats across all seasons, with marked differences in species richness and abundance. Benthic habitats displayed higher overall species richness and invertebrate abundance compared to hyporheic zones, indicating that surface sediments support greater biodiversity. This difference can likely be attributed to DO and food resource availability, and physical habitat heterogeneity in benthic environments, which foster conditions favorable to a diverse and abundant community (Rosi-Marshall and Wallace 2002; Mathers et al. 2017). Also, small pore spaces between sediment particles physically select species with elongated body forms (Marmonier et al. 1993; Descloux et al. 2014). These results align with findings from previous studies by Descloux et al. (2013) and Lencioci and Spitale (2014), who similarly noted that benthic invertebrate communities display higher abundance and species richness compared to hyporheic communities. Further examination of the hyporheic habitats in relation to their proximity to the channel reveals pronounced distinctions among the three sampled locations, with T2 and T3 exhibiting a high degree of similarity, while T1 is intermediate; the tendency of decreasing both abundance and richness in the order of benthic zone, T1, and two other locations was clear. This divergence likely arises from site-specific variations in local conditions within each hyporheic habitat. As Dole-Olivier et al. (2022) emphasize, local hydrological dynamics and substrate conditions are pivotal in shaping hyporheic faunal distribution and community structure within gravel bar environments. However, community composition analyses are necessary to elucidate the community assembly mechanisms behind these patterns.

Our LCBD results demonstrated the role of hyporheic zones in providing habitat for taxonomically unique invertebrate community, despite their relatively low abundance and taxonomic richness. In particular, the pivotal ecological importance of T3 sites in sustaining a distinct invertebrate community was evidenced by consistently high LCBD values across seasons. LCBD highlights sites or samples irrespective of the conservation values of species behind, thus the interpretation with the careful examination of taxa with high SCBD is fundamental (Sor et al. 2018). This uniqueness of T3 was largely driven by hyporheic-dwelling taxa, *Pseudocrangonyx yezonis* and isopod species, and are well adapted to the unique conditions in the hyporheic zones (Boulton et al. 2010; Marin and Palatov 2024). In contrast, T2 and T1 was intermediate in relation to the benthic zone. Together with the understanding of the decrease in abundance and taxonomic richness, LCBD patterns among the locations were largely driven by the occurrence of unique native taxa with low abundance in the hyporheic zone. T1 and T2 hyporheic zones displayed lower LCBD values, suggesting that their invertebrate communities were less unique and potentially more similar to each other or the benthic zone. This could be due to a variety of factors, such as less pronounced hydrological differences between these locations, lower habitat complexity, or a lack of species that are uniquely adapted to hyporheic environments. However, it is also important to note that T2 also exhibited a high abundance of *Pseudocrangonyx yezonis* and isopod species together with occasionally high SCBD scores, underscoring its ecological significance as a critical habitat for unique hyporheic-dwelling species in certain seasonal contexts. This emphasizes the relevance of T2 in supporting invertebrate diversity despite its lower overall contribution to beta diversity compared to T3.

The ecological significance of *Pseudocrangonyx yezonis* and isopod species at T3 extends beyond their role as biodiversity contributors. These organisms perform essential functions that are vital to maintaining the river ecosystem function and stability, thus conserving their habitat is important. They are key contributors to nutrient cycling and sediment processing within hyporheic habitats. They play a central role in the decomposition of organic matter, breaking down detritus and facilitating nutrient release into the ecosystem, which promotes primary productivity and supports higher trophic levels (Fiser et al. 2019; Kralj-Fišer et al. 2020). Additionally, these species contribute to sediment turnover and bioturbation, processes that enhance the exchange of oxygen and nutrients between the surface and subsurface waters, improve sediment structure, and support water filtration functions within the river ecosystem (Nogaro and Burgin 2014; Shrivastava et al. 2021).

These processes are vital for the overall health and function of the hyporheic zone, as it enhances its ability to act as a filter for organic materials and pollutants, promoting water purification and improving habitat quality.

As our study examined two sites, we recognize that the limited site replication imposes constraints on the broader spatial applicability of our findings, and caution should be exercised when generalizing these results to other river systems, particularly given the spatial heterogeneity present at regional or landscape scales. Nevertheless, the incorporation of multiple within-site replications across various habitats (benthic, T1, T2, T3) combined with seasonal collection still allowed us to capture fine-scale spatial and temporal variability in the benthic and hyporheic habitat of gravel bars. This approach enabled us to provide novel insights into the ecological role and importance of the lateral extension of HZ in gravel bars in enhancing the overall riverine invertebrate. While acknowledging the spatial limitations, the results of this study could provide valuable baseline information that may inform future research conducted at broader spatial scales and support comparative assessments aimed at evaluating the generality and applicability of these patterns in other river systems.

### Environmental filtering of hyporheic invertebrate communities

The invertebrate community in T1 shows a strong positive association with dissolved oxygen (DO) levels, with higher oxygen concentrations supporting increased abundance and diversity in this habitat. This relationship suggests the role of DO as a critical factor in shaping hyporheic community dynamics. Invertebrates rely on sufficient oxygen to sustain their metabolic processes, which drive growth, reproduction, overall fitness, and even emergence. Without adequate DO, many invertebrates may experience restricted growth or survival, leading to reduced community resilience and diversity (Galic et al. 2023; Thakur et al. 2023). One of the key reasons for T1’s higher DO levels compared to T2 and T3 across all seasons probably lies in its proximity to the wetted channel. Positioned within the channel itself, T1 consistently receives inflow from the oxygen-rich surface water. This continuous seepage ensures that oxygen is readily available to hyporheic invertebrates, creating a favorable environment that directly supports their metabolic demands (Kaufman et al. 2017). In contrast, T2 and T3 are located further from the wetted channel, reducing the inflow of oxygenated water. As a result, DO levels in T2 and T3 are lower, which may limit the survival of invertebrates with higher oxygen needs and, consequently, contribute to the differences in community structure and diversity observed between these locations. The reduction of DO with an increasing distance from channels is in line with previous studies (Peralta-Maraver et al. 2018).

Despite the lower dissolved oxygen (DO) levels in T2 and T3 compared to T1, these areas exhibit stable thermal conditions, which are likely driven by the higher contribution of groundwater discharge in the hyporheic zone of gravel bars. Groundwater discharge acts as a natural buffer against the rapid temperature fluctuations that typically occur in surface water environments, creating a more consistent thermal regime in the hyporheic zone (Dole-Olivier 2011; Land and Peters 2023). Thermal stability plays a critical role in supporting the ecological processes of hyporheic invertebrates. Temperature influences various biological functions, such as embryonic development, growth, emergence, and metabolism (Beracko and Revajova 2019). For ectothermic organisms like invertebrates, stable temperatures reduce physiological stress and allow these processes to proceed within optimal limits (Bonacina et al. 2022; Kazmi et al. 2022). This stability can provide a crucial refuge from the more dynamic and unpredictable conditions of the wetted channel, where rapid temperature fluctuations can disrupt metabolic balance and hinder survival. The benefits of thermal stability, however, come with the trade-off of lower DO levels. Dissolved oxygen is essential for aerobic metabolism, and low-oxygen environments can limit the activity and distribution of many aquatic taxa (Thakur et al. 2023). Nonetheless, hyporheic invertebrates that inhabit these zones often exhibit specialized adaptations to cope with oxygen stress. These adaptations may include low metabolic oxygen requirements, the ability to use anaerobic pathways temporarily, or behaviors that enable them to access microhabitats with higher oxygen concentrations (Kim et al. 2017; Stubbington et al. 2019). This interplay between thermal stability and DO availability highlights the unique ecological niche provided by T2 and T3. While the reduced oxygen levels may exclude some species, the consistent thermal environment creates a suitable habitat for certain taxon that are well-adapted to hyporheic conditions.

The positive association observed in community structure in T2 and T3 with increasing distance from the wetted channel may be a mere indirect reflection of DO and/or thermal stability as main filters of community structure. However, there are other potentially important causal factors and distance itself can also independently act as a filtering agent. Firstly, as distance from the channel increases, the habitat likely becomes less conducive for invertebrates due to limited space and food resources. Although we did not measure sediment characteristics along with the distance gradient, sediment porosity and thus space may become a strong limiting factor (Mather et al. 2017). Secondly, the supply of food resources is presumably limited in the hyporheic zone away from the main channel because river-produced resources cannot physically reach the area (DelVecchia et al. 2019). Lastly, many insect invertebrates, such as species from Ephemeroptera, Plecoptera, and Trichoptera orders, are reliant on surface water for their emergence as adults. This emergence process requires an upward movement from the hyporheic zone through the sediment to the surface water, followed by the transition out of the water column (Baxter et al. 2017; Dorff and Fin 2020). If individuals venture further into T2 and T3, the increasing distance from the wetted channel complicates their pathway to emergence, creating additional physiological challenges and energetic demands. The taxa inhabiting T2 and T3 are primarily permanent hyporheic dwellers that do not require emergence later in their life cycle. This allows them to persist in these environments despite the constraints imposed by increasing distance from the wetted channel, highlighting their ecological specialization and resilience to hyporheic conditions. In this context, intriguingly, *Lepidostoma* was consistently found in both benthic and hyporheic zones including T2 as also reported earlier (Negishi et al. 2022). The eventual fate of them is unknown and warrants further research.

## Conclusion

This study examines the role of the lateral extension of the hyporheic zone in gravel bars in contributing to total riverine invertebrate diversity. We observed that hyporheic invertebrate community composition is markedly distinct from those in benthic zones, and among the hyporheic habitats, stable gravel bars support a particularly unique community composition. This uniqueness is primarily driven by hyporheic-dwelling taxa, particularly *Pseudocrangonyx yezonis* and isopod species, which are absent from benthic habitat and less prevalent in other hyporheic sites. Lateral gradients of various physicochemical conditions in the hyporheic zone extending from the river channel to the hyporheic zone beneath active and stable gravel bars are the key drivers of spatial turnover of community and overall taxonomic richness. Altered sediment dynamics due to flow regulation have reduced the natural formation of gravel bars and compromised their structural integrity, threatening their ecological function (Sumitomo et al. 2018; Nakamura et al. 2020). Globally, sediment augmentation, involving the deliberate addition of sediments to river systems, and artificial high-flow releases are among common management strategies in addressing ecosystem degradation (Nakamura et al. 2020). Our findings provide an additional empirical foundation to the vital role of such efforts in sustaining and enhancing riverine biodiversity and their functions.

## Acknowledgments

We want to express our gratitude to the Obihiro Regional Office of the Hokkaido Development Bureau, Ministry of Land, Infrastructure, Transport and Tourism (MLIT) for their help and assistance in the field. We also like to express our gratitude to Dr. K. Toyoda for his assistance in doing the water quality analyses and Dr. Y. Hayakawa for his help in doing the UAV photograph of the site.

## Author contributions

MJSV - Conceptualization, methodology, data analysis, writing; JNN: Conceptualization - methodology, data analysis, writing, supervision, field surveys, laboratory analysis; YW - Field surveys, data analysis, review & editing; SY - Field survey & laboratory analysis; JTM: Field surveys, laboratory analysis, data analysis, review & editing.

## Funding

This study was partly supported by research funds for the Tokachi and Ishikari rivers provided by MLIT (18056588) and JSPS KAKENHI (18H03408, 22H03787, and 23H02241).

## Data availability

All of the supporting tables and figures are present in the supplementary information. The data that support the findings of this study is available upon reasonable request from the corresponding author.

## Declarations

### Ethical approval

No ethical violations occurred in this research.

### Conflict of interest

The authors declare that they have no conflict of interest.

## Notes

### Competing Interest Statement

The authors have declared no competing interest.

