## Supplement figures for "Contribution of the lateral extension of the hyporheic zone in gravel bars in shaping river invertebrate diversity"

**Supplementary File 1 (Figures)**

Mc Jervis S. Villaruel[1,3], Junjiro N. Negishi[2], Junyi Wu[1], Shuyu Yao[1], Janine T. Mihara[1]

[1]Division of Environmental Science and Development, Graduate School of Environmental Science, Hokkaido University, Sapporo City, Hokkaido, 060-0810, (corresponding author, M.J.S. Villaruel,)

[2]Faculty of Environmental Earth Science, Hokkaido University, Sapporo City, Hokkaido, 060-0810, Japan

[3]Department of Science and Technology, Research Unit - Philippine Science High School - Main Campus, Diliman Quezon City, Philippines

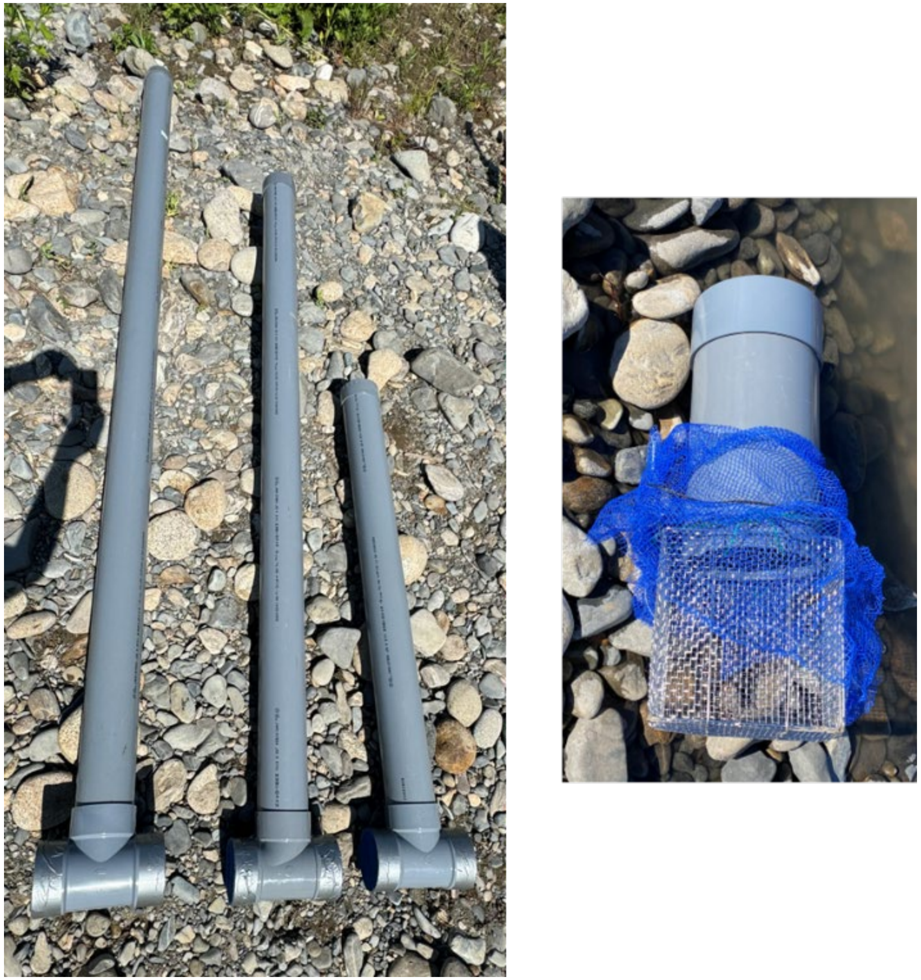

Figure S1. PVC pipes used for hyporheic community collection

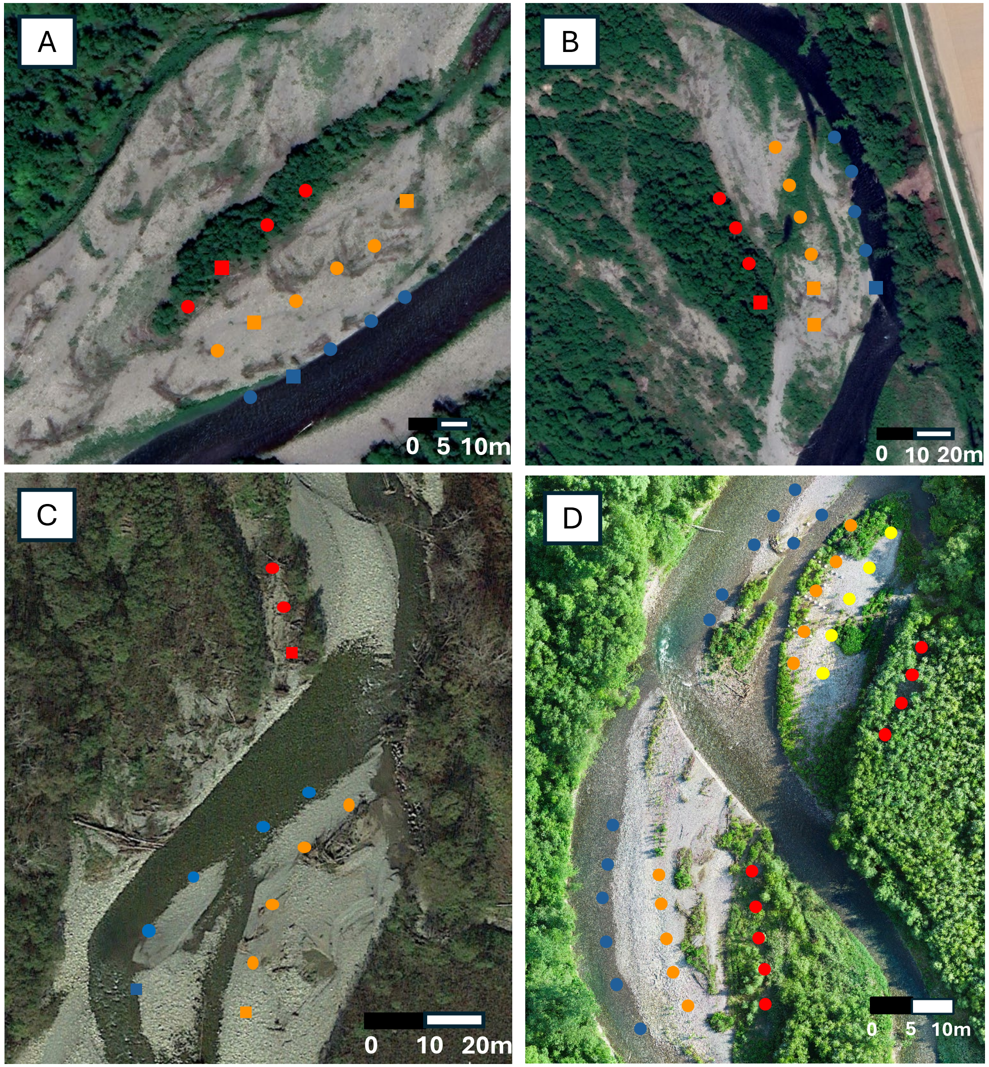


Figure S2. Pipe design used showing the increasing distance of pipe from the wetted channel (blue = wetted channel; orange and yellow = active gravel bar; red = stable gravel bars). (A) L7 site in June 2022 collection, (B) L2 sites in June 2022 collection, (C) L2 in December 2022, March 2023 and May 2023 collection, (D) L2 and Island Bar (IB) in June 2024 collection.

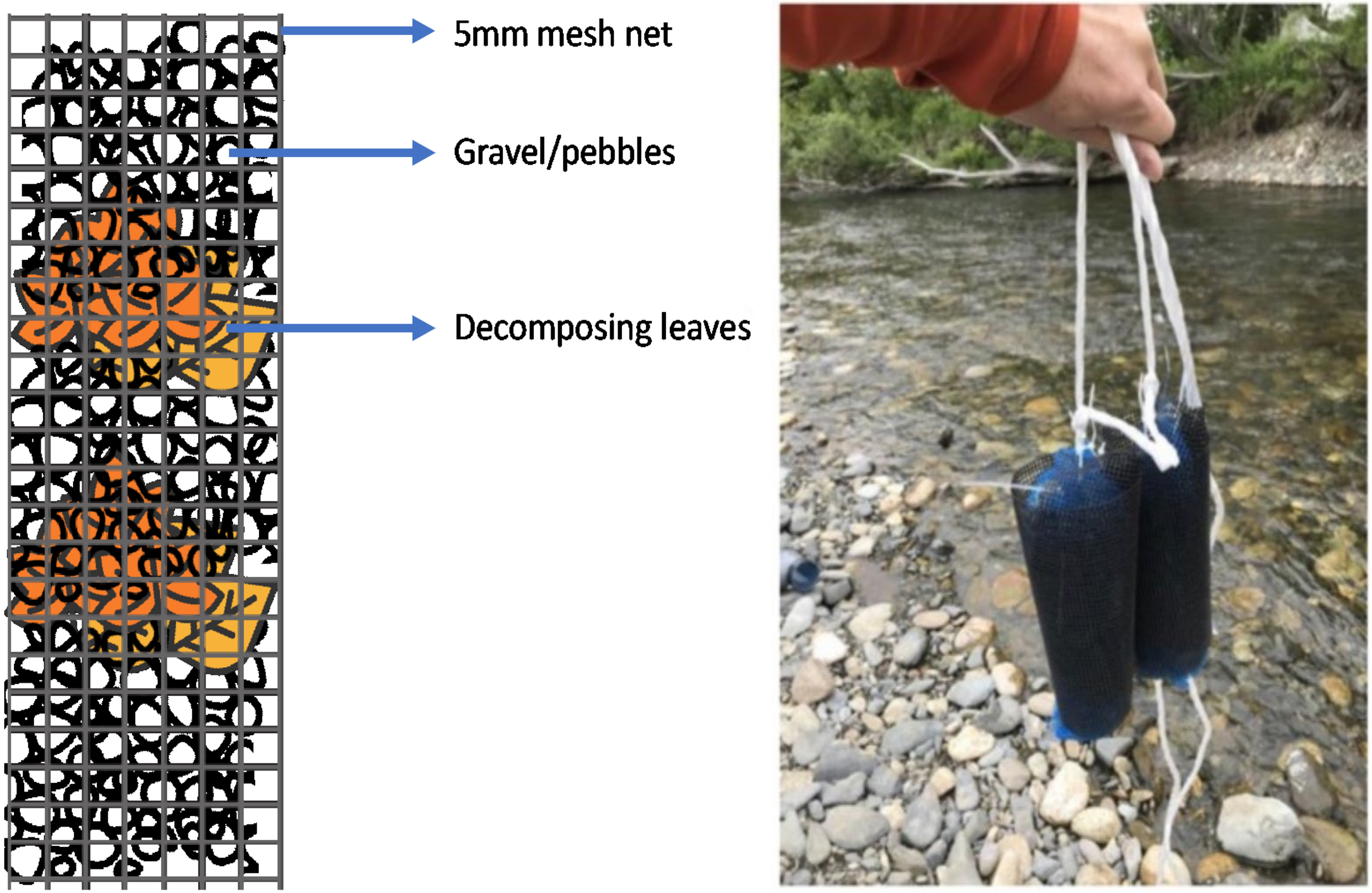


Figure S3. Design and components of hyporheic traps

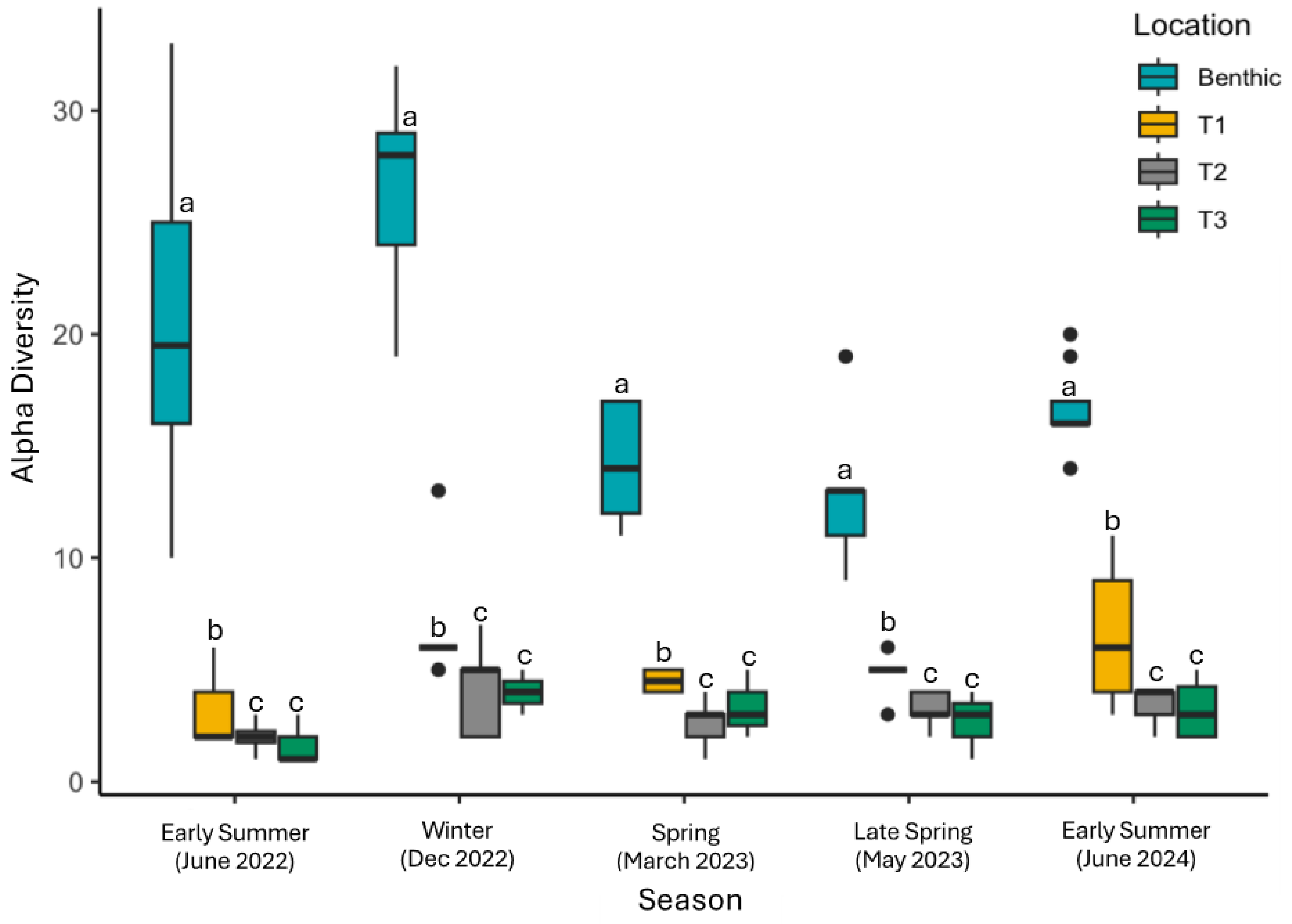


Figure S4. Comparison of alpha diversity across different locations (Benthic, T1, T2, T3) in each season. Alphabetical letters denote results of multiple comparisons based on GLMM, those with the same letters are not statistical different. Note that multiple comparisons were performed among locations only without season factors considered because of the absence of interaction term (see the text).

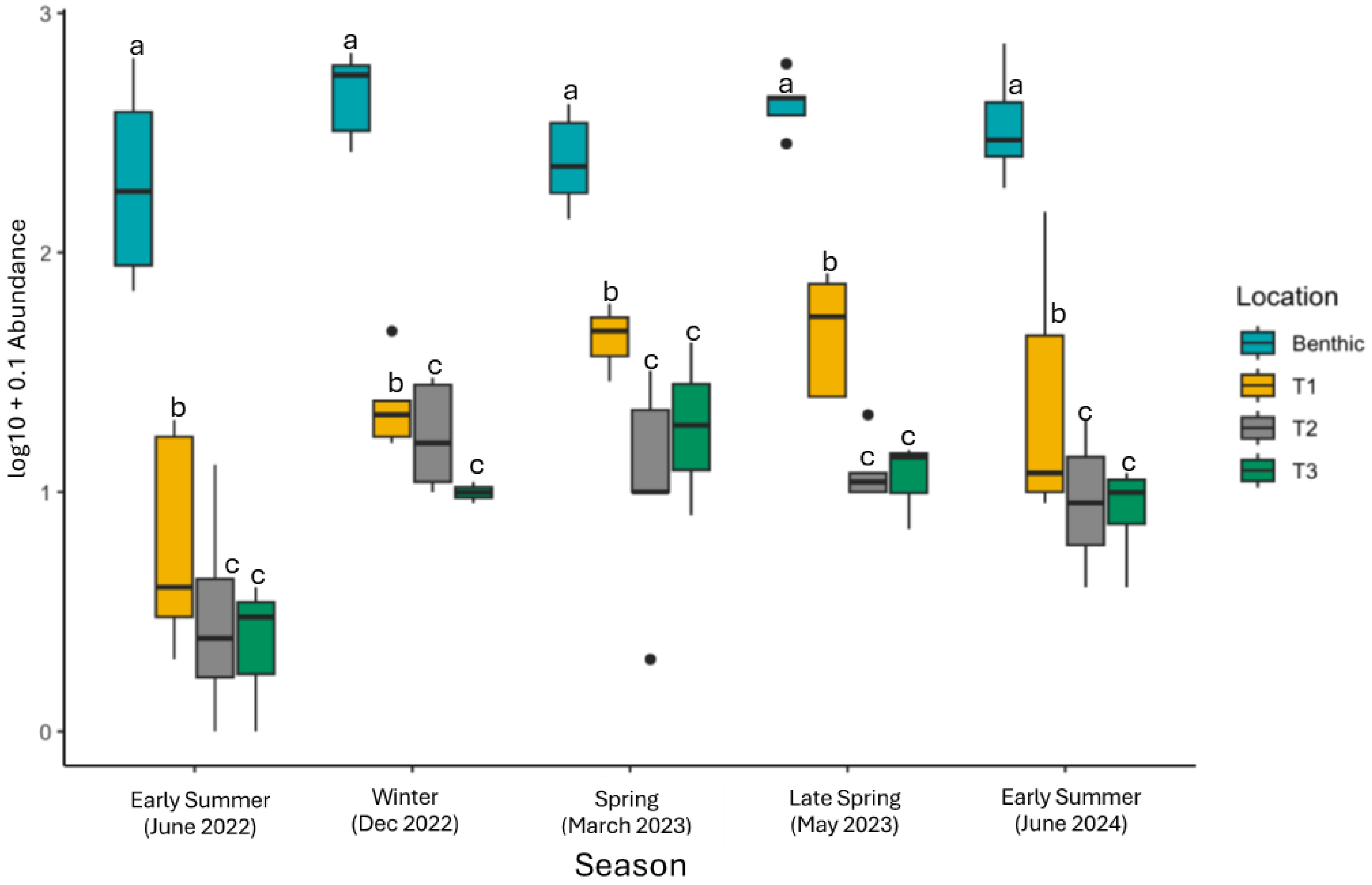


Figure S5. Comparison of invertebrate abundance among different locations across different seasons. Alphabetical letters denote results of multiple comparisons based on GLMM, those with the same letters are not statistical different. Multiple comparisons were performed among locations only without season factors considered because of the absence of interaction term (see the text).

.

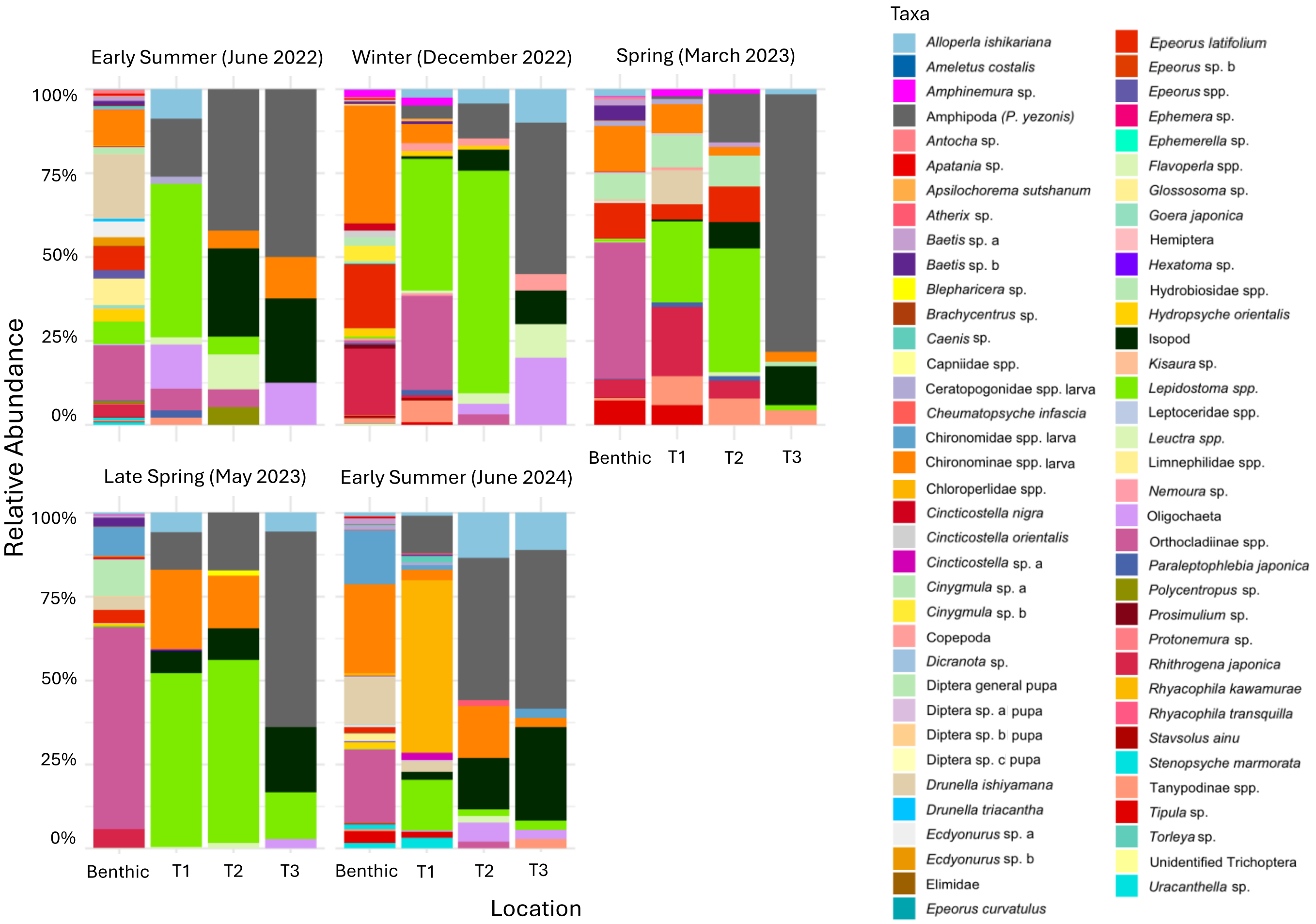


Figure S6.

Composition and relative abundance of invertebrates collected from benthic and different hyporheic locations across different seasons (see the text).
