## Supplement tables for "Contribution of the lateral extension of the hyporheic zone in gravel bars in shaping river invertebrate diversity"

### Supplementary File 2 (Tables)

Mc Jervis S. Villaruel[1,3], Junjiro N. Negishi[2], Junyi Wu[1], Shuyu Yao[1], Janine T. Mihara[1]

[1]Division of Environmental Science and Development, Graduate School of Environmental Science, Hokkaido University, Sapporo City, Hokkaido, 060-0810, (corresponding author, M.J.S. Villaruel,)

[2]Faculty of Environmental Earth Science, Hokkaido University, Sapporo City, Hokkaido, 060-0810, Japan

[3]Department of Science and Technology, Research Unit - Philippine Science High School - Main Campus, Diliman Quezon City, Philippines

Table S1. Summary of sample count collected for different locations during the study period.

| Season | Site | Location | Pipe Number | Number of Samples |
|---|---|---|---|---|
| Early Summer (June 2022) | L7 | Benthic | N/A | 5 |
| | | T1 | 5 | 3 |
| | | T2 | 5 | 2 |
| | | T3 | 5 | 1 |
| | L2 | Benthic | N/A | 5 |
| | | T1 | 5 | 2 |
| | | T2 | 5 | 2 |
| | | T3 | 5 | 2 |
| Winter (Dec 2022) | L2 | Benthic | N/A | 5 |
| | | T1 | 5 | 5 |
| | | T2 | 5 | 5 |
| | | T3 | 3 | 3 |
| Spring (March 2023) | L2 | Benthic | N/A | 5 |
| | | T1 | 5 | 4 |
| | | T2 | 5 | 5 |
| | | T3 | 3 | 3 |
| Late Spring (May 2023) | L2 | Benthic | N/A | 5 |
| | | T1 | 5 | 5 |
| | | T2 | 5 | 5 |
| | | T3 | 3 | 3 |
| Early Summer (June 2024) | L2 | Benthic | N/A | 5 |
| | | T1 | 5 | 2 |
| | | T2 | 5 | 2 |
| | | T3 | 4 | 2 |
| | IB | Benthic | N/A | 5 |
| | | T1 | 6 | 3 |
| | | T2 | 5 | 3 |
| | | T3 | 5 | 3 |

Table S2. Environmental parameters measured in the benthic and hyporheic locations across different locations.

| Season | Location | EC (mS/m) | Temperature ($^{O}C$) | pH | D.O. (mg/L) | D.O. saturation | TN (mg/L) | TP (mg/L) | Cl (mg/L) | $NO_3$ (mg/L) | $SO_4$ (mg/L) | Velocity (cm/s) | Depth (cm) |
|---|---|---|---|---|---|---|---|---|---|---|---|---|---|
| Early Summer (June 2022) | Benthic | 3.61±0.08 | 17.51±0.29 | 6.85±0.11 | 8.43±0.09 | 94.90±3.10 | 0.22±0.03 | 0.03±0.01 | 1.15±0.03 | 1.55±0.01 | 3.46±0.04 | 64.4±8.79 | 25.8±2.26 |
| | T1 | 3.88±0.24 | 16.08±1.11 | 6.65±0.07 | 7.73±0.12 | 83.81±0.14 | 0.20±0.04 | 0.04±0.01 | 1.21±0.10 | 1.53±0.05 | 3.46±0.19 | N/A | N/A |
| | T2 | 3.88±0.22 | 15.75±0.76 | 6.64±0.10 | 7.47±0.21 | 82.3±0.15 | 0.19±0.05 | 0.05±0.01 | 1.24±0.13 | 1.43±0.11 | 3.58±0.19 | N/A | N/A |
| | T3 | 3.94±0.27 | 15.30±0.47 | 6.58±0.04 | 7.30±0.15 | 81.97±0.13 | 0.15±0.03 | 0.06±0.01 | 1.16±0.09 | 1.51±0.06 | 3.42±0.08 | N/A | N/A |
| Winter (Dec 2022) | Benthic | 3.92±0.07 | 4.21±0.03 | 6.14±0.02 | 11.46±0.04 | 94.49±8.12 | 0.74±0.08 | 0.02±0.01 | 1.46±0.06 | 2.38±0.20 | 3.73±0.14 | 87.1±3.56 | 34.3±1.96 |
| | T1 | 4.68±0.03 | 3.58±0.20 | 6.21±0.04 | 11.60±0.08 | 88.10±0.11 | 0.79±0.11 | 0.08±0.03 | 1.71±0.03 | 3.05±0.05 | 4.14±0.06 | N/A | N/A |
| | T2 | 4.64±0.02 | 2.44±0.14 | 6.40±0.04 | 11.74±0.06 | 87.80±2.10 | 0.46±0.10 | 0.03±0.01 | 1.42±0.36 | 2.37±0.59 | 3.39±0.85 | N/A | N/A |
| | T3 | 4.88±0.41 | 2.64±0.22 | 6.34±0.17 | 11.22±0.21 | 84.5±3.06 | 0.74±0.06 | 0.03±0.01 | 1.26±0.13 | 2.30±0.19 | 3.47±0.32 | N/A | N/A |
| Spring (March 2023) | Benthic | 6.16±0.20 | 7.10±0.73 | 6.78±0.10 | 11.17±0.21 | 92.80±0.01 | 1.73±0.43 | 0.01±0.00 | 2.04±0.10 | 2.11±0.23 | 5.35±0.16 | 99.6±5.34 | 37.7±2.32 |
| | T1 | 6.30±0.02 | 8.48±0.60 | 6.97±0.11 | 10.64±0.08 | 85.6±4.09 | 2.36±0.09 | 0.20±0.06 | 2.39±0.04 | 2.19±0.05 | 6.07±0.05 | N/A | N/A |
| | T2 | 6.41±0.08 | 7.06±0.32 | 6.56±0.11 | 10.33±0.17 | 84.42±6.18 | 2.40±0.08 | 0.14±0.02 | 2.85±0.26 | 2.57±0.05 | 5.98±0.05 | N/A | N/A |
| | T3 | 6.72±0.25 | 5.23±0.67 | 6.70±0.08 | 10.82±0.15 | 84.66±2.22 | 2.66±0.22 | 0.21±0.01 | 2.33±0.07 | 3.77±0.84 | 6.38±0.31 | N/A | N/A |
| Late Spring (May 2023) | Benthic | 3.60±0.57 | 17.58±2.36 | 6.12±1.01 | 9.12±1.50 | 97.4±0.20 | 0.32±0.02 | 0.01±0.00 | 1.32±0.21 | 2.07±0.20 | 3.75±0.57 | 75.2±3.01 | 28.7±3.84 |
| | T1 | 3.48±0.02 | 16.22±0.56 | 6.73±0.14 | 9.11±0.04 | 95.10±4.87 | 0.35±0.87 | 0.03±0.01 | 1.41±0.04 | 2.08±0.06 | 3.58±0.05 | N/A | N/A |
| | T2 | 3.56±0.03 | 15.77±0.39 | 6.89±0.14 | 8.74±0.08 | 89.05±5.76 | 0.43±0.06 | 0.03±0.01 | 1.43±0.04 | 2.11±0.05 | 3.61±0.07 | N/A | N/A |
| | T3 | 3.42±0.03 | 14.80±0.27 | 6.20±0.19 | 8.99±0.31 | 89.20±5.6 | 0.34±0.06 | 0.01±0.00 | 1.46±0.03 | 2.17±0.10 | 3.60±0.03 | N/A | N/A |
| Early Summer (June 2024) | Benthic | 5.61±0.19 | 16.60±0.13 | 6.53±0.17 | 8.08±0.01 | 98±0.11 | 0.81±0.01 | 0.03±0.01 | 1.14±0.01 | 1.79±0.01 | 3.90±0.01 | 26.4±9.28 | 20.1±1.67 |
| | T1 | 6.40±0.37 | 15.24±0.28 | 6.20±0.38 | 8.05±0.37 | 88.9±9.19 | 1.42±0.09 | 0.02±0.01 | 1.13±0.27 | 1.46±0.64 | 3.48±0.12 | N/A | N/A |
| | T2 | 5.87±0.07 | 15.22±0.34 | 6.32±0.24 | 7.85±0.11 | 77.1±7.18 | 0.65±0.18 | 0.03±0.01 | 1.95±0.39 | 1.53±0.60 | 3.41±0.73 | N/A | N/A |
| | T3 | 6.13±0.22 | 14.62±0.25 | 6.21±0.17 | 7.22±0.33 | 73.1±5.23 | 0.95±0.23 | 0.16±0.10 | 1.87±0.35 | 1.44±0.59 | 3.63±0.09 | N/A | N/A |

Table S3. Results of GLMMs testing the effects of location and season and their interactions on alpha diversity of invertebrates. Full model and reduced model were compared using log-likelihood tests. When the p-value of the first 1st reduced model was insignificant, it was compared to 2nd reduced models. Superscripts in the p-values indicate the variables excluded from the model for comparison with the reduced model. P-values in bold font denote statistical significance.

| Alpha Diversity | logLik | AIC | p-value |
|---|---|---|---|
| Full model | | | |
| Season (S), Location (L), S x L | -210.03 | 462.06 | 0.2475 |
| 1st Reduced Model | | | |
| S, L | -217.47 | 452.95 | **<0.0001**$^{S}$; **<0.0001**$^{\mathbf{L}}$ |
| 2nd Reduced Model | | | |
| S | -232.68 | 475.37 | |
| L | -495.08 | 1002.16 | |

Table S4. Results of GLMMs testing the effects of location and season and their interactions on abundance of invertebrates. Full model and reduced model were compared using log-likelihood tests. When the p-value of the first 1st reduced model was insignificant, it was compared to 2nd reduced models. Superscripts in the p-values indicate the variables excluded from the model for comparison with the reduced model. P-values in bold font denote statistical significance.

| Alpha Diversity | logLik | AIC | p-value |
|---|---|---|---|
| Full model | | | |
| Season (S), Location (L), S x L | -26.085 | 96.171 | 0.1812 |
| $1^{st}$ Reduced Model | | | |
| S, L | -34.197 | 88.394 | **<0.0001$^{S}$; <0.0001$^{L}$** |
| $2^{nd}$ Reduced Model | | | |
| S | -50.211 | 112.423 | |
| L | -114.223 | 196.171 | |

Table S5. Results of GLMMs testing the effects of location and season and their interactions on LCBD values. Full model and reduced model were compared using log-likelihood tests. When the p-value of the first 1st reduced model was insignificant, it was compared to 2nd reduced models. Superscripts in the p-values indicate the variables excluded from the model for comparison with the reduced model. P-values in bold font denote statistical significance.

| LCBD Values | logLik | AIC | p-value |
|---|---|---|---|
| Full model | | | |
| Season (S), Location (L), T x L | 94.592 | -145.18 | 0.6313 |
| 1st Reduced Model | | | |
| S, L | 89.679 | -159.36 | **<0.05**$^{S}$; **<0.0001**$^{L}$ |
| 2nd Reduced Model | | | |
| S | 88.473 | -164.95 | |
| L | 71.345 | -128.69 | |

Table S6. Results of the Redundance Analysis and Permutational ANOVA of the environmental parameters and the hyporheic invertebrate community composition. Number of permutations: 999.

| Model RDA Hyporheic Invertebrate Community and Environmental Parameters | | | | |
|---|---|---|---|---|
| | df | Variance | F | Pr (> F) |
| Model | 12 | 0.19073 | 2.7339 | **0.001***** |
| Residual | 51 | 0.44814 | | |
| *Significant codes:* | *0 '****' 0.001 '***' 0.01 '**' 0.05 '*'* | | | |

Table S7. Results of Permutational Analysis of Variance (PERMANOVA) testing the difference in centroids of the group in RDA multivariate space [A] and Pairwise PERMANOVA results comparing the different groups in the RDA multivariate space [B] using RDA scores.

| A. PERMANOVA | $R^2$ | F | p-value |
|---|---|---|---|
| Location | 0.129 | 4.427 | **0.004**** |
| Residual | 0.871 | | |
| Total | 1.000 | | |

| B. Pairwise PERMANOVA | $R^2$ | F | p-value |
|---|---|---|---|
| T1 vs T2 | 0.009 | 0.445 | **0.010*** |
| T1 vs T3 | 0.187 | 8.554 | **0.002**** |
| T2 vs T3 | 0.129 | 5.463 | 0.654 |

*Significant codes: 0 '****' 0.001 '***' 0.01 '**' 0.05 '**
